# Regulation of Hippo signaling by Atrophin in the developing *Drosophila* wing

**DOI:** 10.1101/2025.10.09.681328

**Authors:** Deimante Mikalauskaite, Cordelia Rauskolb, Tom Lehan, Srividya Venkatramanan, Lilia Carnes, Kenneth D. Irvine

## Abstract

Organ development is directed through integration of signaling networks and their transcriptional programs. We have investigated connections between Hippo signaling and the transcriptional co-repressor Atrophin. We find that Atrophin modulates Hippo signaling outputs in the developing *Drosophila* wing and does so in distinct ways in different regions. Near the dorsal-ventral boundary, loss of Atrophin leads to upregulation of targets of the Hippo pathway transcription factor Yorkie. This is explained by impairment of Notch signaling, and consequent downregulation of Vestigial, which normally competes with Yorkie for binding to Scalloped. In proximal regions of the wing disc, loss of Atrophin leads to downregulation of Yorkie activity. This is explained by downregulation of Dachs, as Dachs inhibits Warts, the central kinase controlling Yorkie activity. Downregulation of Dachs is explained by modulation of its upstream regulators Dachsous and Four-jointed, which is explained in turn by our discovery that Atrophin interacts genetically and physically with Vestigial and competes with Scalloped for Vestigial binding. These studies define new roles for Atrophin and enhance our understanding of the interplay of transcriptional activators and repressors that modulate Hippo signaling to shape wing development.

## Introduction

Transcriptional regulatory networks orchestrate cell fate decisions and behaviors by integrating cell signaling with intrinsic genetic programs. The Hippo signaling network regulates cell fate and organ growth largely through inhibiting the transcriptional co-activator protein Yorkie (Yki) (Misra and Irvine, 2018; Zhong et al., 2024). Yki is regulated by Hippo signaling through the kinase Warts (Wts), which phosphorylates Yki to promote its cytoplasmic localization (Dong et al., 2007; Huang et al., 2005; Oh and Irvine, 2008).

Conversely, unphosphorylated Yki accumulates in the nucleus where it partners with the DNA binding protein Scalloped (Sd) to promote transcription of target genes (Goulev et al., 2008; Wu et al., 2008; Zhang et al., 2008). Sd also has additional partners that it can interact with to direct alternative transcriptional programs (Fig. 1A). In the absence of Yki, Sd interacts with the transcriptional co-repressor Tondu-domain-containing Growth Inhibitor (Tgi) to repress target genes (Koontz et al., 2013). Hippo signaling activity thus switches Sd from participating in activation of target genes, with Yki, to repression of target genes, with Tgi. Sd also can associate with E2F1 (Zhang et al., 2017), which leads to downregulation of Yki-Sd target genes, or with Vestigial (Vg), to activate genes involved in wing development (Halder et al., 1998; Paumard-Rigal et al., 1998; Simmonds et al., 1998). Yki can also interact with other transcription factors, including Homothorax, Mad, and Trithorax-like (Trl, also called GAF) (Bayarmagnai et al., 2012; Oh and Irvine, 2011; Oh et al., 2013; Peng et al., 2009). Here, we describe investigations of a link between the transcriptional co-repressor Atrophin (Atro, encoded by *grunge*) and Hippo signaling.

**Figure 1.**
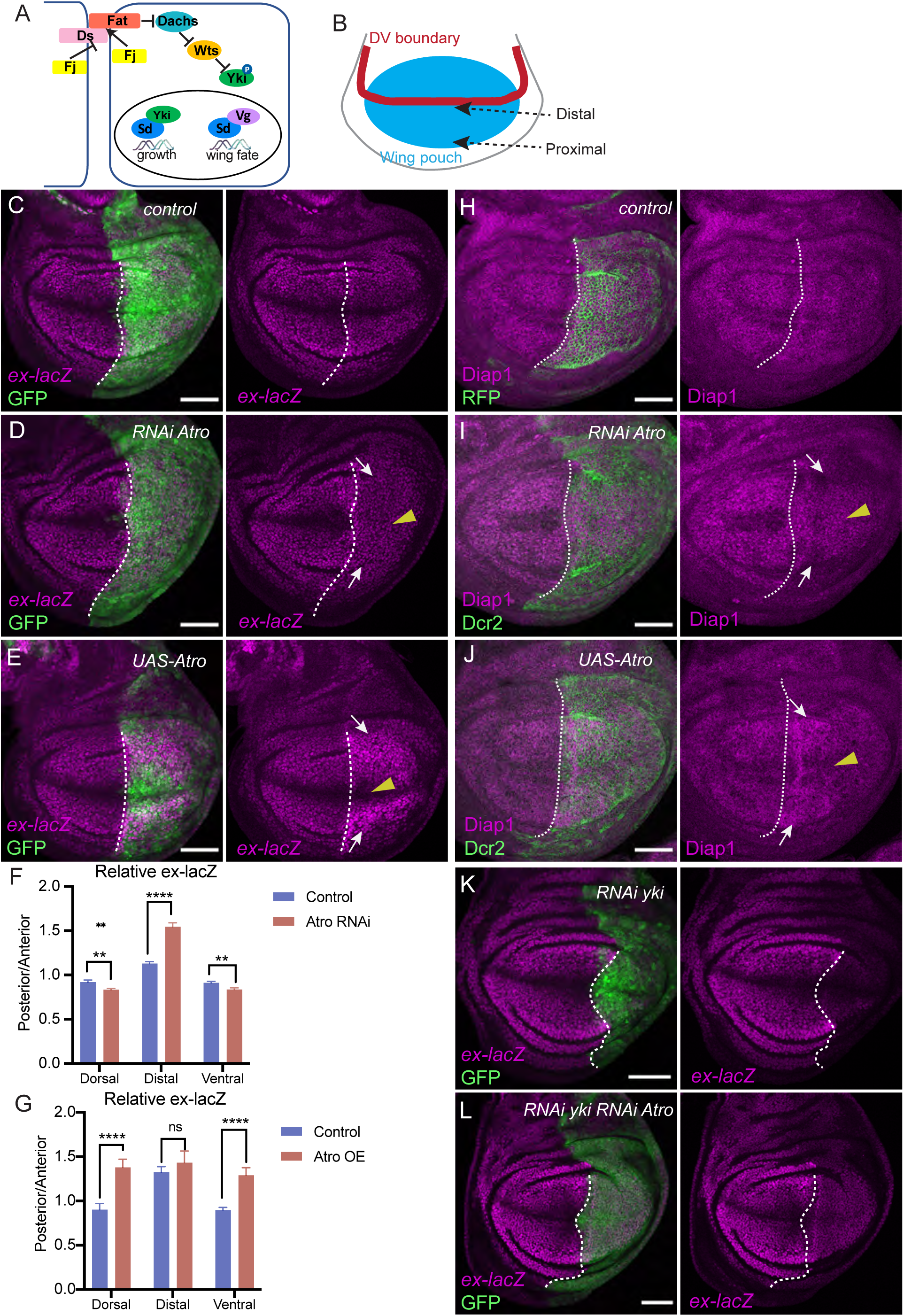
Regulation of Yki target genes by Atrophin. (A) Simplified schematic of Ds-Fat branch of Hippo pathway, showing repression of Yki via phosphorylation by Wts. Unphosphorylated Yki can enter the nucleus, where it partners with Sd to promote transcription of genes increasing growth. Alternatively, Sd can partner with Vg to promote expression of genes specifying wing cell fate. B) Schematic of wing region of the wing disc, showing location of developing wing (wing pouch) and DV boundary. The center of the wing pouch is distal, and peripheral regions are proximal. (C-E) Expression of *ex-lacZ* (magenta) in third instar wing discs from *en-Gal4 UAS-GFP ex-lacZ; tub-Gal80^ts^ UAS-Dcr2* crossed to control (Oregon-R, OR) (C), *UAS-RNAi-Atro [HMS00756]* (D), or UAS-3x:FLAG-Atro (E). Posterior compartment is labeled by GFP expression (green). (F-G) Quantification of relative ex-lacZ in posterior versus anterior cells of wing discs from animals of genotype described in C-E in distal regions (near DV boundary) and proximal regions (Dorsal and Ventral). N=11 for *UAS-RNAi-Atro* and control animals (F), N=8 for *UAS-3xFLAG:Atro* and 7 for control (G). Significance of differences calculated by unpaired Student’s t-test. (H-J) Expression of Diap1 (magenta) in third instar wing discs from *en-Gal4 UAS-RFP; tub-Gal80^ts^ UAS-dcr2* crossed to control (OR) (H), *en-Gal4; tub-Gal80^ts^ UAS-Dcr2* crossed to *UAS-RNAi-Atro* (I), or *UAS-Atro III* (J). Posterior compartment is labeled by RFP (H) or Dcr2 (I,J) (green). (K,L) Expression of *ex-lacZ* (magenta) in third instar wing discs from *en-Gal4 UAS-GFP ex-lacZ; tub-Gal80^ts^ UAS-Dcr2* crossed to *UAS-RNA-yki[HMS00041]* (K), *UAS-RNA-yki[HMS00041] UAS-RNAi-Atro[KK100675]* (L). Posterior compartment is labeled by GFP expression (green). Dashed lines mark anterior-posterior compartment boundary, yellow arrowheads point to the D-V boundary region, white arrows point to proximal wing pouch. Scale bars=50 μm.

Atro is an evolutionarily conserved protein that plays important roles at multiple stages of *Drosophila* development (Charroux et al., 2006; Erkner et al., 2002; Wang and Tsai, 2008; Yeung et al., 2017; Zhang et al., 2013), and in mutant form is linked to human neurodegenerative disease (Koide et al., 1994; Nagafuchi et al., 1994). Published observations hint at potential connections between Atro and Hippo signaling. Clones of cells mutant for Atro in the eye imaginal disc upregulate expression of *four-jointed (fj)* (Fanto et al., 2003), which is also a Yki target gene. Atro was also reported to be able to associate with the protocadherin Fat, which is an upstream regulator of Hippo signaling (Fanto et al., 2003). Atro can also associate with, and co-localize on chromatin with, Trl (Yeung et al., 2017), which is also a co-factor for Yki (Oh et al., 2013). These observations prompted us to directly examine potential functional connections between Atro and Hippo signaling, using the developing *Drosophila* wing as a model.

The *Drosophila* wing has been extensively employed as a model for investigations of organ growth, patterning and morphogenesis (Tripathi and Irvine, 2022). It forms from the wing imaginal disc, a cluster of undifferentiated cells that undergo extensive growth and patterning during larval development. The wing imaginal disc also gives rise to the wing hinge, and notum of the adult fly; the region of the disc that will give rise to the wing, termed the wing pouch (Fig. 1B), is specified by Vg, acting in conjunction with Sd (Halder et al., 1998; Paumard-Rigal et al., 1998; Simmonds et al., 1998; Williams et al., 1991). In addition to specifying wing fate, and promoting wing cell survival and proliferation, Vg also contributes to wing patterning, as Vg is expressed in a gradient with highest levels of expression near the dorsal-ventral (DV) boundary, and lower levels away from the DV boundary, in more proximal regions (Cho and Irvine, 2004; Kim et al., 1996; Williams et al., 1993; Zecca et al., 1996). This gradient of Vg expression contributes to regulation of wing growth and patterning in part through regulation of key components of Ds-Fat signaling (Cho and Irvine, 2004; Zecca and Struhl, 2010).

Ds-Fat signaling is mediated by intercellular interaction of two large protocadherins, Ds and Fat (Fulford and McNeill, 2020; Gridnev and Misra, 2022). Binding between Ds and Fat is modulated by the Golgi-localized kinase Fj, which phosphorylates Ds and Fat cadherin domains (Brittle et al., 2010; Ishikawa et al., 2008; Simon et al., 2010). In the developing wing, Fj is positively regulated by Vg and expressed in a similar DV-high to proximal-low gradient (Cho and Irvine, 2004; Zecca and Struhl, 2010). Conversely, Ds is repressed by Vg and expressed in a proximal-high to D-V-low gradient (Clark et al., 1995; Rogulja et al., 2008; Zecca and Struhl, 2010). These gradients of Ds and Fj bias interactions between Ds and Fat, such that Ds and Fat become polarized within cells, with Ft concentrated on the proximal sides of cells and Ds on the distal sides of cells (Ambegaonkar et al., 2012; Brittle et al., 2012). Downstream signaling is mediated principally through Fat regulation of the myosin family protein Dachs (Fig. 1A) (Cho et al., 2006; Mao et al., 2006). Membrane localization of Dachs is repressed by Fat, and consequently Dachs localization is also polarized to the distal sides of cells in the developing wing. Dachs then mediates downstream effects of Ds-Fat signaling on Hippo signaling, planar cell polarity (PCP), and morphogenesis (Fulford and McNeill, 2020; Gridnev and Misra, 2022; Strutt and Strutt, 2021).

We report here that Atro regulates the expression of Yki target genes in the developing wing imaginal disc. This regulation is complex, reflecting Atro’s connection to multiple factors. Near the DV boundary, we find that Atro downregulates Yki target gene expression, likely through an essential contribution to Notch signaling, which modulates Yki activity here. Layered over this localized downregulation of Yki target genes is a broader upregulation of Yki target genes. This broader upregulation is mediated through regulation of Hippo signaling, as revealed by effects of Atro on nuclear localization of Yki. Our investigations further define a regulatory pathway connecting Atro to Yki regulation, involving upregulation of Ds levels and downregulation of Dachs levels. We describe a potential mechanism for Atro’s effects on Ds expression, through identification of a competition between Atro and Sd for physical association with Vg. These studies identify Atrophin as a modulator of Hippo signaling and contribute to our understanding of the genetic regulatory network that controls Hippo signaling to instruct wing development.

## Results

### Atrophin regulates the expression of Hippo pathway target genes in wing discs

To investigate the possibility that Atro regulates Hippo signaling, we examined the consequences of Atro knockdown in wing discs. Atro protein is detected in nuclei throughout the wing disc as revealed by staining with anti-Atro antibody, or by visualizing Atrophin expressed from a genomic Atrophin: GFP transgene (Fig. S1A,B). We knocked down Atro in posterior cells by expressing either of two different *UAS-RNAi* transgenes under the control of *en-Gal4*, which both anti-Atro staining and examination of Atrophin:GFP confirmed were highly effective at reducing Atro protein levels (Fig. S1C-E).

We then examined the influence of Atro knock down on expression of *expanded* (*ex*), a well-established direct target of Yki, using the *ex* transcriptional reporter *ex-lacZ* (Hamaratoglu et al., 2006). Depletion of Atro had complex effects on *ex-lacZ* expression. In the central region of the developing wing pouch, near the DV boundary, where *ex-lacZ* levels are normally low, *ex-lacZ* levels were increased by Atro RNAi (Fig. 1C,D,F).

Conversely, in the most proximal regions of the wing pouch, where *ex-lacZ* levels are normally higher, *ex-lacZ* levels were reduced by Atro RNAi (Fig. 1D,F). Together, these effects led to a flattening of the normal gradient of *ex-lacZ* expression in the wing pouch. Similar effects were observed using an independent Atro RNAi line (Fig. S1F).

To determine whether other Hippo pathway target genes were also affected by knockdown of Atro, we examined expression of another direct target gene, *Diap1*, using anti-Diap1 antibody staining. Diap1 was affected similarly to *ex-lacZ*, with elevated expression in cells near the DV boundary, and reduced expression in the proximal wing pouch (Fig. 1H,I).

To determine whether expression of Hippo pathway target genes could also be altered by increased Atro expression, we used UAS-Atro transgenes to over-express Atro in posterior cells under *en-Gal4* control. This maintained reduced expression of *ex* and *Diap1* near the DV boundary, while increasing their expression in the proximal wing pouch (Fig. 1E,G,J). Thus, both increasing and decreasing Atro levels altered the expression of Hippo pathway target genes, but distinct effects were observed in distal versus proximal regions of the developing wing.

### Atro partially suppresses the effects of altered Yki activity

To begin to investigate the mechanism by which Atro affects Yki target gene expression, we examined genetic interactions between *yki* and *Atro*. When Yki levels were reduced in posterior wing cells by RNAi, the expression of Yki target genes was strongly reduced throughout the posterior compartment (Fig. 1K, Fig. S1G). When knockdown of Atro was combined with knockdown of Yki, the effect of Yki knockdown was partially reversed, as low levels of *ex-lacZ* and Diap1 were observed throughout the wing pouch (Fig. 1L, Fig. S1H). To increase Yki activity, we over-expressed an activated form of Yki (Yki^S168A^) throughout posterior cells, which increased *ex-lacZ* and Diap1 expression (Fig. S1I,K).

When Yki^S168A^ expression was combined with Atro over-expression, expression of both *ex-lacZ* and Diap1 was reduced near the center of the wing pouch, but otherwise appeared similar to that observed in wing discs over-expressing Yki^S168A^ (Fig. S1J,L). The observation that alterations in Atro levels can partially reverse the consequences of loss or gain of Yki in the distal wing raised the possibility that Atro might act in parallel to Yki to regulate Yki target genes in this region.

### Atro promotes Notch activity at the DV boundary

As Atro differentially affected Hippo pathway target genes in different regions of the wing disc, we hypothesized that it could be acting through distinct targets in different regions. The DV boundary of the wing disc is specified by Notch activation, raising the possibility that the differential effects observed there could be due to effects on Notch signaling.

Indeed, RNAi-mediated down-regulation of Atro has previously been associated with reduced expression of Notch target genes along the DV boundary of the wing disc, including *wingless* and *cut* (Wang et al., 2018; Yeung et al., 2017). It was further suggested that this is due to reduced transcription of Notch, based on reported analysis of a *Notch-lacZ* reporter (Wang et al., 2018). However, we note that the expression pattern reported for *Notch-lacZ* (Wang et al., 2018) does not match that reported for Notch protein (Fehon et al., 1991). Moreover, when we used anti-Notch antibodies to examine Notch in wing discs with Atro-knock down in posterior cells, we did not observe any effects on Notch protein levels or distribution (Fig. 2A).

**Figure 2.**
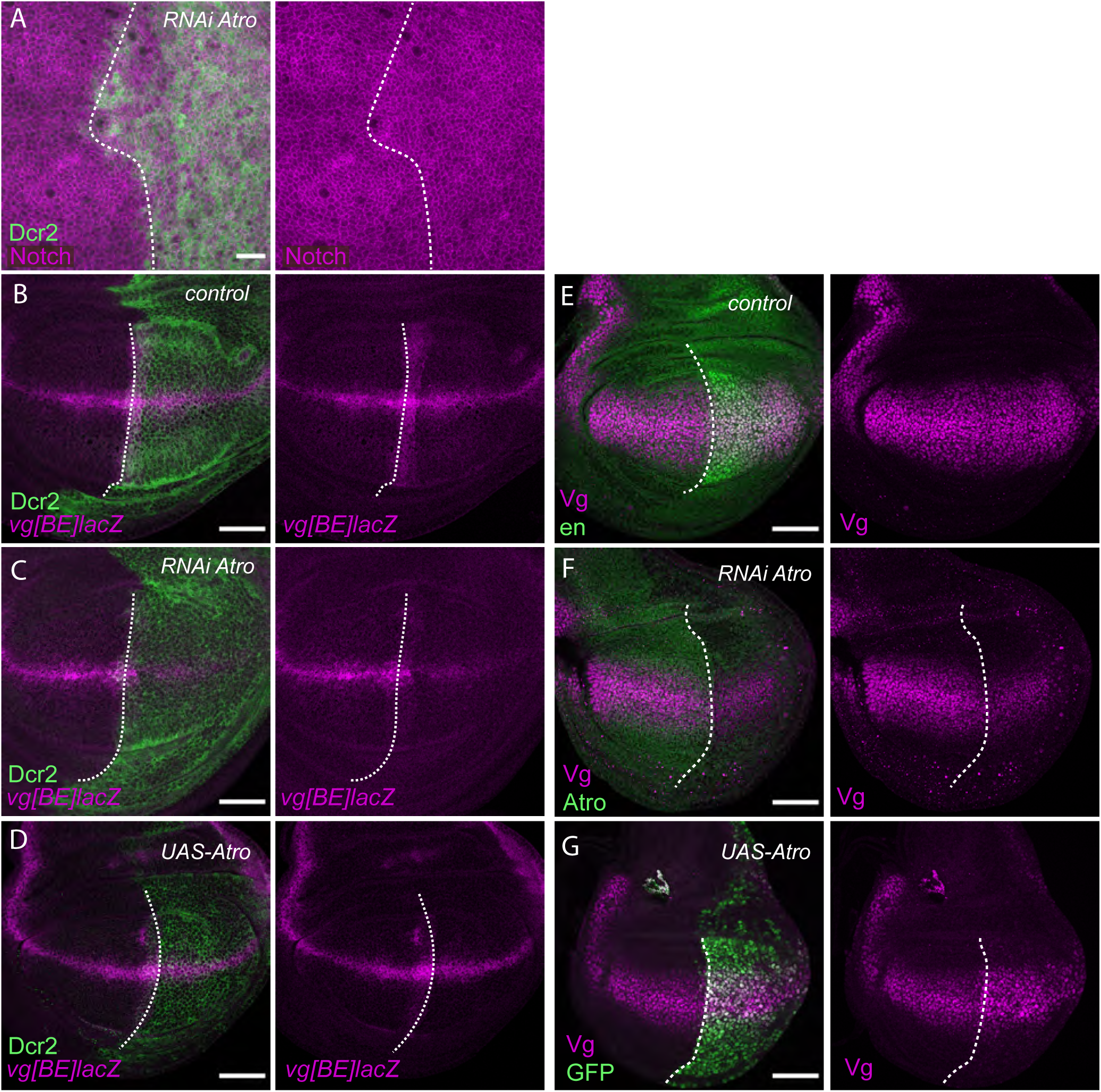
Atrophin controls signaling of the Notch pathway. (A) Notch protein staining (magenta) in wing disc expressing *en-Gal4 tub-Gal80^ts^ UAS-Dcr2* crossed to *UAS-RNAi-Atro*. Posterior compartment is labeled by Dcr2 (green), scale bar=10 um. (B-D) Expression of *vg[BE]-lacZ* (magenta) in wing discs from *en-Gal4 tub-Gal80^ts^ UAS-Dcr2* crossed to *vg-[BE]-lacZ* (B), *UAS-RNAi-Atro[HMS00756] vg-[BE]-lacZ* (C), and *UAS-Atro vg-[BE]-lacZ* (D). Posterior compartment is labeled by Dcr2 (green). (E-G) Expression of Vg protein (magenta) in wing discs from *en-Gal4 tub-Gal80^ts^* crossed to O-R (control) with posterior compartment marked by En stain (green) (E), *UAS-RNAi-Atro* with posterior compartment marked by loss of Atro antibody staining (green), or *UAS-Atro UAS-GFP* with posterior compartment labeled by GFP expression (green). Scale bars=50 μm. Dashed lines mark anterior-posterior compartment boundary.

Although the mechanism by which Atro affects Notch signaling output remains uncertain, it could nonetheless potentially explain the effect of Atro on Hippo pathway targets near the DV boundary, as it has been reported that Notch inhibits Yki activity in wing discs (Djiane et al., 2014). This inhibition was ascribed to competition between Vg and Yki for their shared DNA binding partner, Scalloped (Sd). Notch promotes Vg expression along the DV boundary through the *vg* boundary enhancer, and elevated Vg could suppress Yki target gene expression by out-competing Yki for association with Sd. Consistent with this possibility, when we analyzed expression driven by the *vg* boundary enhancer using a lacZ reporter (*vg-[BE]lacZ)* that is directly activated by Notch signaling (Kim et al., 1996), we observed that *vg-[BE]lacZ* levels were reduced in posterior cells when Atro levels were knocked down by RNAi (Fig. 2B,C). Vg protein levels were also visibly reduced by Atro RNAi (Fig. 2E,F). Over-expression of Atro, conversely, increased Vg protein staining, although *vg-[BE]lacZ* was not visibly affected (Fig. 2D,G). Thus, altogether our results are consistent with the possibility that downregulation of Vg expression, mediated at least in part through decreased Notch signaling, contributes to the up regulation of Yki target genes near the DV boundary of the wing disc when Atro expression is reduced, and similarly that upregulation of Vg could contribute to the downregulation of Yki targets when Atro is over-expressed.

### Atrophin influences the subcellular localization of Yorkie

While modulation of Vg levels through Notch could explain the effects of Atro on Yki target genes near the DV boundary, distinct mechanisms must be involved in the effects observed in more proximal regions. Two very different types of mechanisms can be envisioned: Atro might influence Yki activity through effects on Hippo signaling, or it might act independently of Hippo signaling, converging on shared targets. To help distinguish between these possibilities, we examined the subcellular distribution of Yki. Knockdown of Atro in posterior cells led to a slight, but statistically significant, reduction in nuclear Yki (Fig. 3A-C). Conversely, over-expression of Atro in posterior cells led to a slight increase in nuclear Yki (Fig. 3D-F). In contrast to the more localized effects on Hippo pathway target genes, these effects of altered Atro levels were observed in both distal and proximal cells. These observations suggest that the positive influence of Atro on expression of Yki target genes in proximal cells is due to an influence on Hippo signaling, which controls nuclear localization of Yki. Conversely, the negative influence of Atro on expression of Yki target genes in distal cells is independent of effects on Hippo signaling.

**Figure 3.**
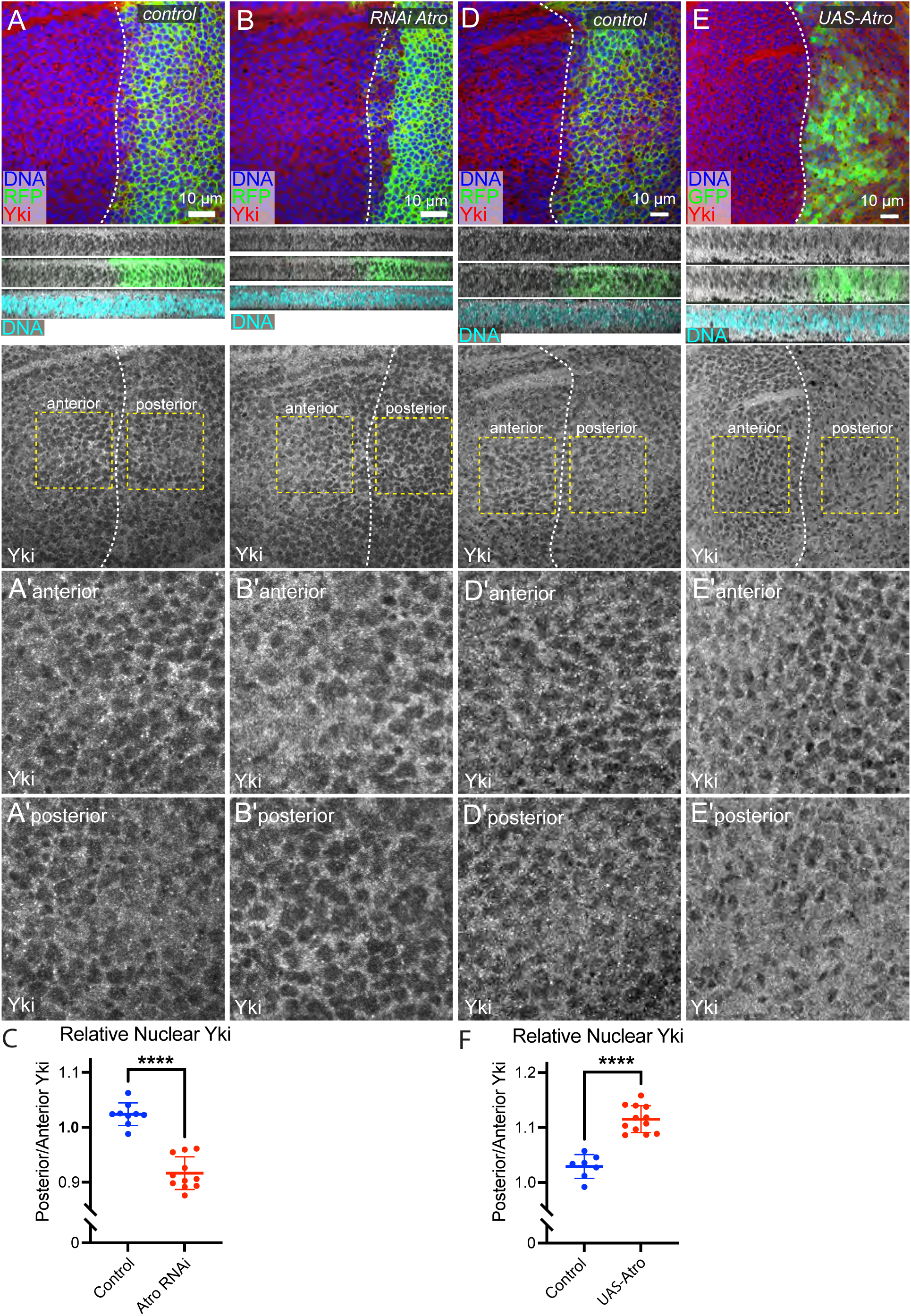
Atrophin promotes Yki nuclear localization. (A,B,D,E) Yki protein staining (red/white) in wing discs expressing *en-Gal4 UAS-RFP(or GFP) tub-Gal80^ts^ UAS-Dcr2* crossed to control (OR) (A), *UAS-RNAi-Atro[HMS00756]* (B), control (OR) (D), or *UAS-3xFLAG:Atro* (E), with posterior compartment marked by expression of RFP or GFP (green) and nuclei marked by DNA staining (blue/cyan). Scale bars=10 μm. Images below show vertical sections through the discs, including Yki, Yki and RFP/GFP, Yki and DNA stain. Below this Yki stain alone is shown in the horizontal section, with dashed yellow boxes outlining anterior and posterior regions shown at higher magnification in the panels further below, identified by prime symbols and anterior or posterior location. (C) Quantification of relative nuclear Yki in posterior vs. anterior compartment cells in control (n=9) and Atro knock down (n=11) wing discs. (F) Quantification of relative nuclear Yki in posterior vs. anterior compartment cells in control (n=7) and Atro over-expression (n=12) wing discs.

### Atrophin regulates the Hippo pathway through Dachsous-Fat signaling

We considered the possibility that Atro might regulate Yki localization by regulating the expression of one or more components of Hippo signaling. ChIP-seq analysis performed in *Drosophila* Schneider line 2 (S2) cells (Yeung et al., 2017) identified prominent Atro peaks at two core components of the Hippo pathway, *wts* and *kibra*. Thus, we examined whether loss of Atro altered expression of *wts* and *kibra* using *wts-lacZ* and *kibra-lacZ* reporters.

When Atro was knocked down in posterior cells under en-Gal4 control, *wts-lacZ* expression was similar or slightly lower than in control cells (Fig. S2A-B). This cannot explain the reduced nuclear Yki observed, because to the extent that a reduction in Wts levels occurs, it would be expected to lead to increased rather than decreased nuclear Yki. Similarly, depletion of Atro driven by *hh-Gal4* also resulted in slightly reduced *kibra-lacZ* levels (Fig. S2C-D). As Wts and Kibra are downstream targets of Yki (Genevet et al., 2010; Park et al., 2016; Sun et al., 2015), the apparent reduction in their expression is likely a consequence of reduced Yki activity.

Hippo signaling is regulated by multiple upstream inputs, and we also investigated whether modulation of these could contribute to Atro’s effects on Hippo signaling. One key upstream input in the wing disc is cytoskeletal tension, which inhibits Wts activity through recruitment of the Ajuba LIM protein (Jub) to adherens junctions (Rauskolb et al., 2014).

When Atro was knocked down in posterior cells, Jub levels at junctions, visualized using a genomic Jub:GFP, were slightly reduced, but this was matched by a decrease in E-cad levels, such that when we quantified the Jub/E-cad ratio, no significant difference between anterior and posterior cells was observed (Fig. S2E-G). We also did not observe a significant effect on Jub:GFP when over-expressing Atro (Fig. S2H-J). These results suggest that altered cytoskeletal tension and Jub recruitment is unlikely to explain the influence of Atro on Yki activity.

Another key upstream regulator of Hippo signaling in wing discs is the Dachsous-Fat pathway (Ds-Ft) (Fulford and McNeill, 2020; Gridnev and Misra, 2022; Strutt and Strutt, 2021). Ds-Ft signaling affects the levels and activity of Wts and Ex through regulating the levels and localization of the myosin family protein Dachs (Bennett and Harvey, 2006; Cho et al., 2006; Feng and Irvine, 2007; Fulford et al., 2023; Mao et al., 2006; Silva et al., 2006; Vrabioiu and Struhl, 2015; Willecke et al., 2006). Using a tagged Dachs:GFP genomic transgene (Bosveld et al., 2012) as a readout of Ds-Fat pathway activity, we found that knockdown of Atro is associated with reduced levels of Dachs at apical junctions (Fig. 4A,B). Some small, discrete puncta of Dachs remain when Atro is knocked down, but overall levels of Dachs are reduced, as confirmed by quantitation of junctional Dachs levels (Fig. 4E). As Dachs normally promotes Yki activity, this reduction in Dachs could in principle account for the reductions in nuclear Yki levels and Yki target gene expression observed when Atro is knocked down. When Atro was overexpressed levels of Dachs appeared increased on some cell junctions (Fig. 4C,D), but quantitation revealed that overall junctional levels of Dachs were actually slightly decreased (Fig. 4F). This suggests that regulation of Dachs could explain decreased Yki activity when Atro is knocked down but not increased Yki activity when Atro is overexpressed.

**Figure 4.**
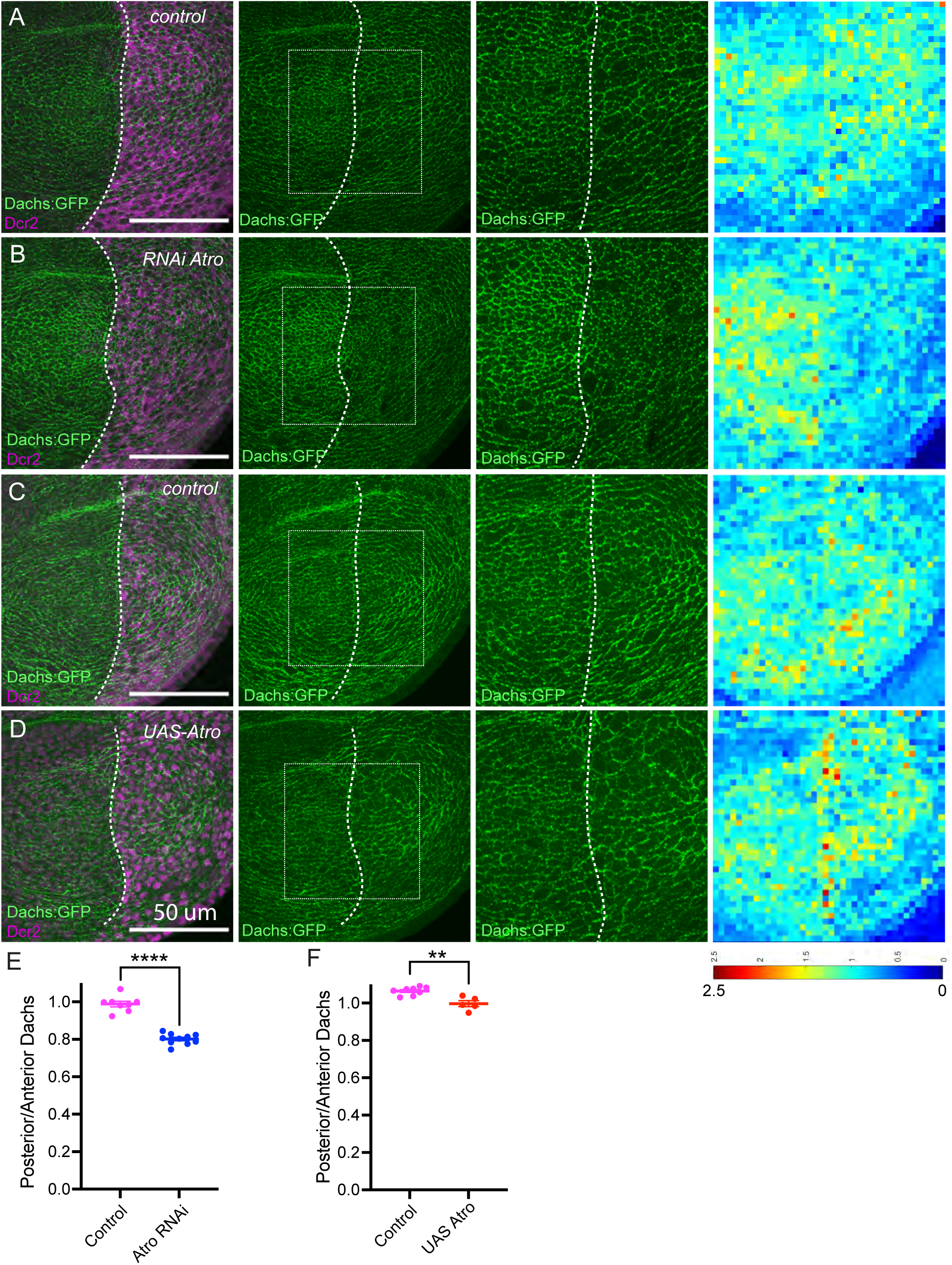
Atrophin regulates Dachs protein levels. (A,B) Dachs:GFP (green) in wing discs expressing *en-Gal4 tub-Gal80^ts^ UAS-Dcr2 Dachs:GFP Ubi-baz:mCherry* crossed to control (OR) (A) or *UAS-RNAi-Atro[HMS00756]* (B). Posterior compartment is labeled by Dcr2 (magenta). (C,D) Dachs:GFP (green) in wing discs expressing *en-Gal4 tub-Gal80^ts^ UAS-Dcr2 Dachs:GFP Ubi-baz:mCherry* crossed to control (OR) (D) or *UAS-3xFLAG:Atro* (E). Posterior compartments are labeled in magenta and are stained with anti-FLAG antibody *(UAS-3XFLAG-Atro)*. For A-D, dashed lines mark anterior-posterior compartment boundary, Dachs:GFP alone is shown to right of dual color image, boxed regions correspond to the magnified images to the right of this, and heat maps at far right show quantitation of Dachs:GFP signal intensity measured as junctional signal intensity (Baz:mCherry). Scale bars=50 μm, scale bar for heat map at bottom. (E) Quantification of relative (normalized to Ecad or Baz:mCherry) junctional Dachs:GFP levels in posterior versus anterior cells for control (n=8) and Atro RNAi (n=10) wing discs of genotypes in A,B, with significance indicated by t-test. (F) Quantification of relative (normalized to Ecad or Baz:mCherry) junctional Dachs:GFP levels in posterior versus anterior cells for control (n= 8) and Atro over-expression (n=5) wing discs of genotypes in C,D, with significance indicated by t test.

To investigate how reduced Atro decreases Dachs levels at junctions, we examined the expression of two key upstream regulators of Ds-Fat signaling, Ds, and Fj. Using a genomic GFP-tagged Ds (Brittle et al., 2012), we observed that Ds levels were substantially upregulated in the wing pouch by knockdown of Atro either throughout the posterior compartment, or in random clones of cells (Fig. 5A-C). As Ds promotes Ft activity, which removes Dachs from cell membranes (Mao et al., 2006; Rogulja et al., 2008), this increased Ds might explain the down regulation of Dachs levels and Yki activity. Indeed, expression of Ds throughout posterior cells was previously reported to reduce expression of Diap1 (Rogulja et al., 2008), and we observed that *ex-lacZ* levels in proximal cells were also reduced by expression of UAS-ds under en-Gal4 control (Fig. 5D,E). When we examined the consequences of directly over-expressing Ds under UAS-Gal4 control, we found that rather than being broadly lower, we observed a distribution suggestive of spontaneous polarization, with some cell junctions having elevated Dachs while others had lower Dachs (Fig S3A). This difference might be explained by differences in the levels of Ds induced by the UAS-Gal4 system versus loss of Atro. Alternatively, it might be explained by additional effects of Atro RNAi. Indeed, we observed that levels of *fj*, as visualized by a *fj-lacZ* reporter, were reduced by Atro RNAi while over-expression resulted in mild elevation of *fj-lacZ* (Fig. S3B-D). Reduced expression of Fj is also associated with reduced Yki activity (Cho and Irvine, 2004; Rogulja et al., 2008). Thus, the altered Dachs levels observed in Atro knockdown cells likely reflect the combined effects of elevated Ds and reduced Fj on Dachs localization.

**Figure 5.**
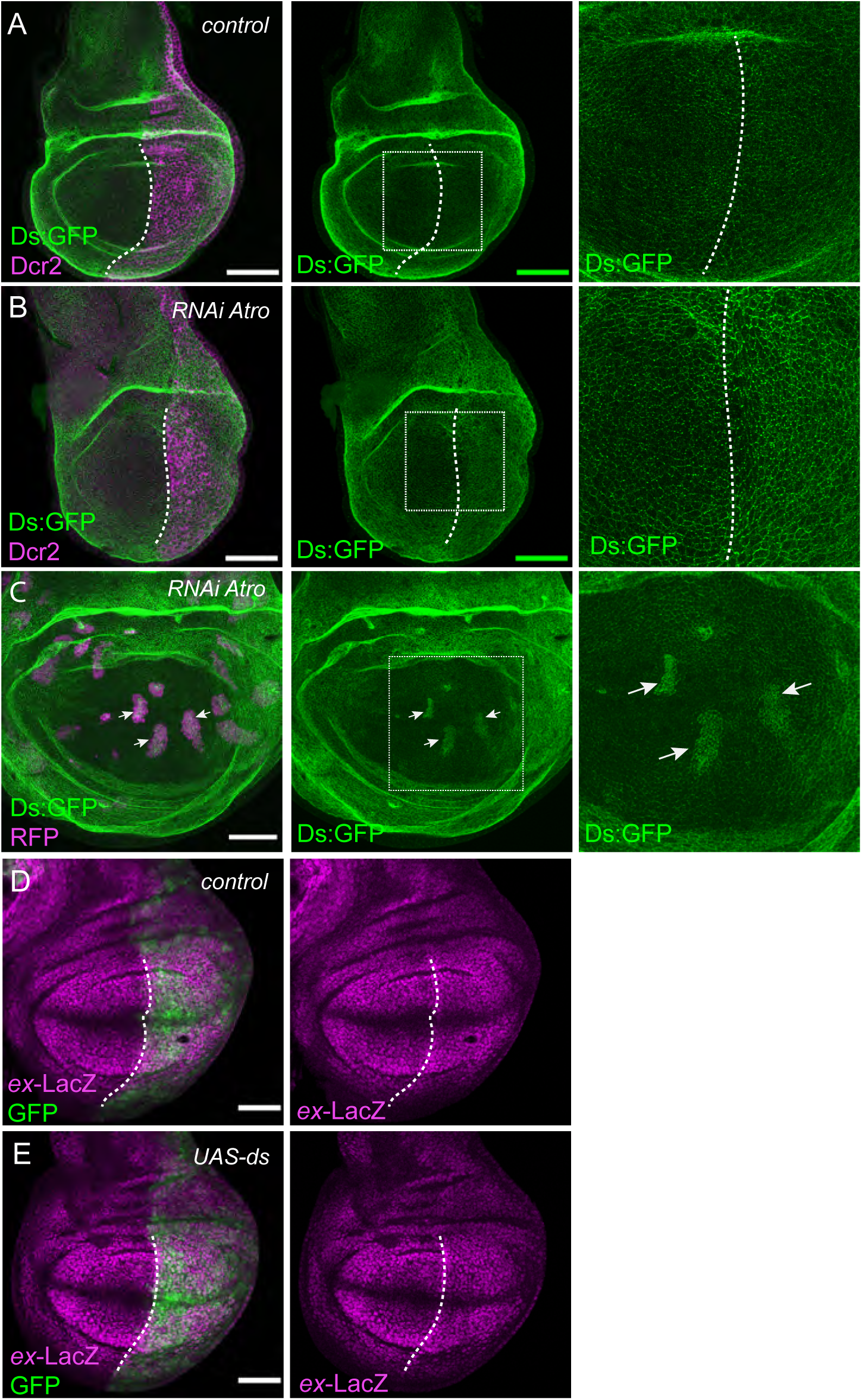
Atrophin controls Dachsous levels. (A,B) Ds:GFP in wing discs expressing *hh-Gal4 tub-Gal80^ts^ UAS-Dcr2 Ds:GFP* crossed to control (OR) (A), *UAS-RNAi-Atro[HMS00756]* (B). Posterior compartments are labeled by Dcr2 (magenta). (C) Ds:GFP in wing disc expressing *act>CD2>Gal4 UAS-nRFP* crossed to *hs-FLP*; *UAS-RNAi-Atro[KK100675]* after heatshock induced expression of FLP created clones expressing *act-Gal4*, labelled by expression of RFP (magenta). Arrows point to examples of clones with elevated Ds:GFP. (D,E) Expression of *ex-lacZ* in wing discs expressing *en-Gal4 UAS-GFP ex-lacZ tub-Gal80^ts^ UAS-Dcr2* crossed to control (OR) (D) or *UAS-ds* (E). Posterior compartment is labeled by UAS-GFP (green). Dashed lines mark anterior-posterior compartment boundary, boxed regions are shown at higher magnification to the right, scale bars=50 μm.

### Atrophin is required for Vg to repress Ds

The complementary expression patterns of Fj and Ds in the developing wing are established in part through the action of Vg, which promotes *fj* expression and inhibits *ds* expression (Cho and Irvine, 2004; Zecca and Struhl, 2010). Vg is a transcriptional co-activator, which partners with Sd to activate downstream target genes (Halder et al., 1998; Paumard-Rigal et al., 1998; Simmonds et al., 1998). How Vg represses *ds* expression is unknown, but the observation that both Vg and Atro normally repress *ds* expression led us to investigate potential functional connections between them, and to consider the possibility that Atro might enable Vg to repress downstream genes like *ds*. Over-expression of Vg, either in random clones or in posterior cells reduced Ds levels in proximal cells (Fig. 6A,B,D), consistent with prior studies (Zecca and Struhl, 2010). When we combined Vg over-expression with Atro knockdown, we observed the Atro knockdown phenotype of Ds upregulation, rather than the Vg over-expression phenotype of Ds downregulation (Figs. 5B,C, 6B-E). The suppression of Vg-mediated Ds repression by Atro knockdown suggests that Atro is required for Vg to repress Ds.

**Figure 6.**
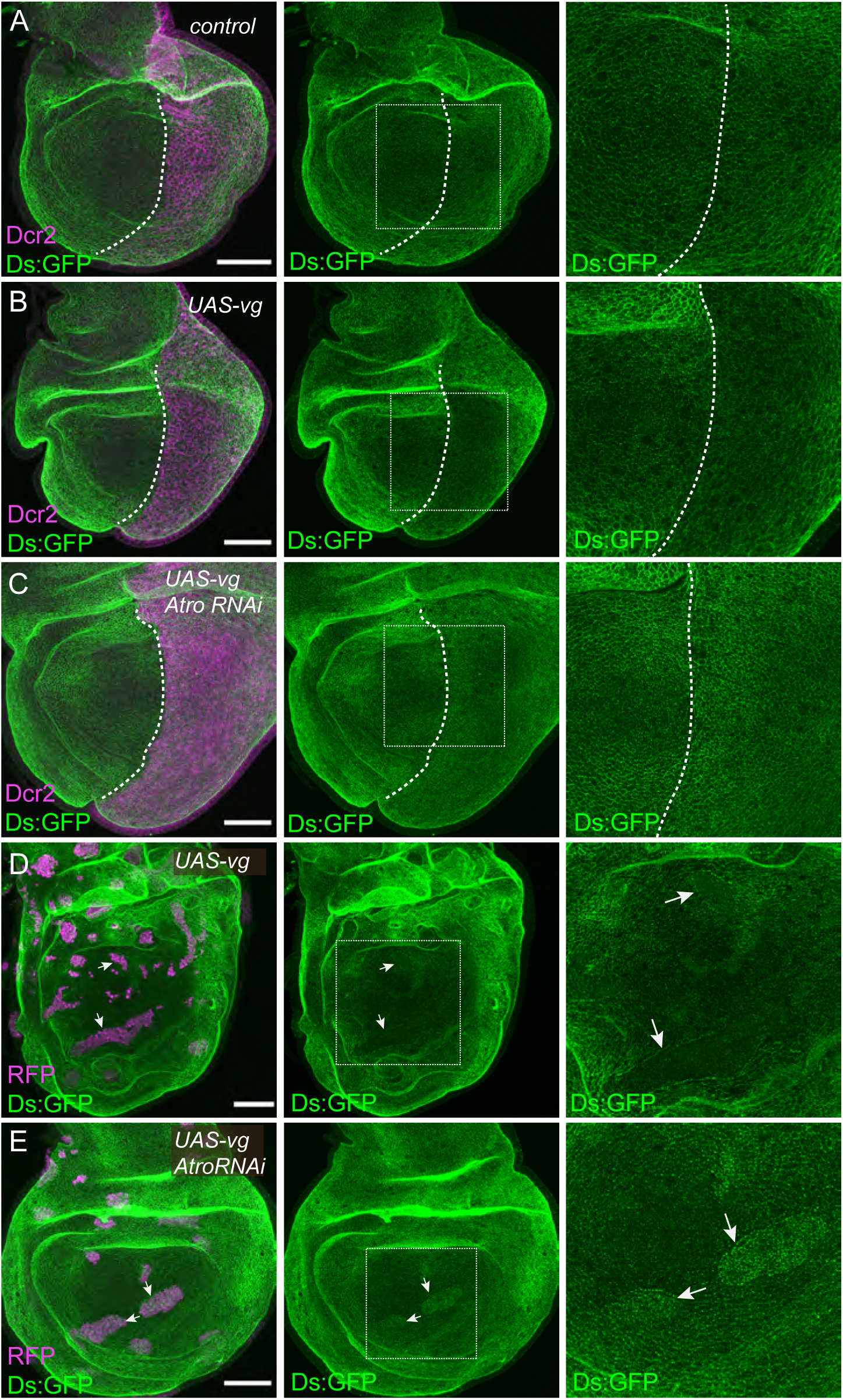
Atrophin is required for repression of Dachsous by Vestigial. (A-C) Ds:GFP in wing discs expressing *hh-Gal4 tub-Gal80^ts^ UAS-Dcr2 Ds:GFP* crossed to control (OR) (A), *UAS-vg* (B), or *UAS-vg UAS-RNAi-Atro[KK100675]* (C). Posterior compartment is labeled by Dcr2 (magenta). (D-E) Ds:GFP in wing discs expressing *act>CD2>Gal4 UAS-nRFP* crossed to *UAS-vg* (D) and *UAS-vg and UAS-RNAi-Atro[KK100675]* after heat shock induced expression of FLP created clones expressing *act-Gal4*, labelled by expression of RFP (magenta). Arrows point to examples of clones with altered Ds:GFP. Dashed lines in A-C mark anterior-posterior compartment boundary, boxed regions are shown at higher magnification to the right, scale bars=50 μm.

### Atrophin physically associates with Vestigial and inhibits Vg-Sd binding

To investigate whether the genetic interaction between Atro and Vg could involve physical association, we co-expressed Atro and Vg in S2 cells and performed co-immunoprecipitation experiments. This revealed that Vg co-precipitated with Atro, indicating that they are together in the same complex, possibly due to direct physical interaction between them (Fig. 7A). As Vg forms a complex with Sd to regulate gene expression, we examined whether Atro could also associate with Sd. However, no co-immunoprecipitation of Sd and Atro was detected (Fig. 7B). We also examined potential complex formation amongst all three proteins. Interestingly, when Atro, Vg, and Sd were co-expressed in S2 cells, Vg no longer co-precipitated with Atro, but instead Vg co-precipitated with Sd (Fig. 7B). This suggested that Atro and Sd could be competing for binding to Vg. To further investigate this, we assayed association between Atro and Vg in the presence of varying amounts of Sd expression. At low Sd expression, some Atro-Vg association could be detected, but as Sd levels increased, Atro-Vg association decreased (Fig. 7B).

**Figure 7.**
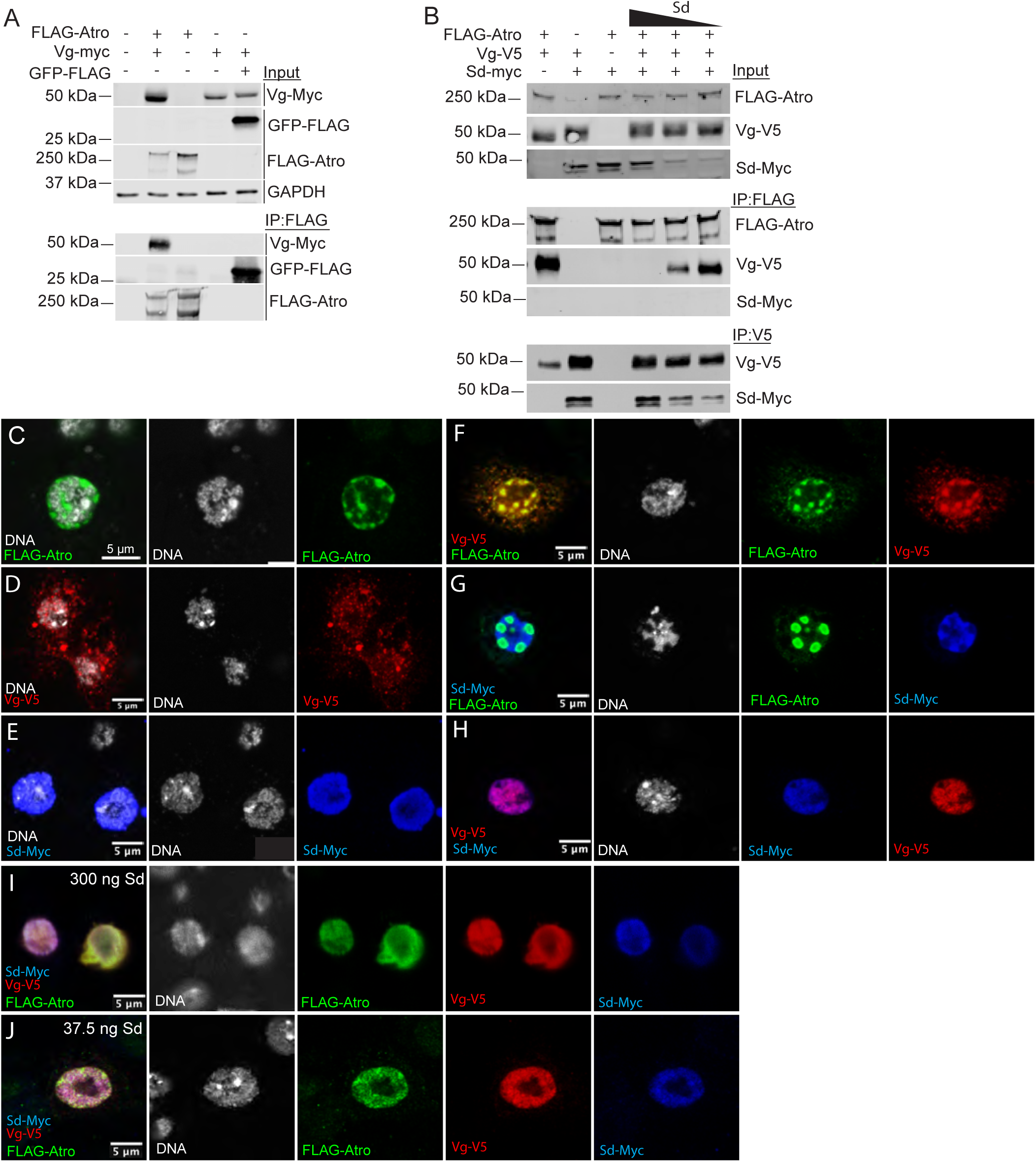
Atrophin binds Vestigial and inhibits Vestigial-Scalloped interaction. (A-B) Western blots showing results of co-immunoprecipitation experiments on lysates of S2 cells transfected with plasmids expressing the proteins indicated at top. Proteins detected (using anti-FLAG, -Myc, -V5, or -GAPDH antibodies) indicated at right of blots, location of protein size standards indicated at left. In (A), top 4 panels (Input) show blots on cell lysates, bottom 3 panels (IP:FLAG) show blots on proteins precipitated by anti-FLAG beads. In (B), three lanes at far right have decreasing amounts of Sd expression plasmid transfected (300, 75, and 37.5 ng), top 3 panels (Input) show blots on cell lysates, next 3 panels (IP:FLAG) show blots on proteins precipitated by anti-FLAG beads, bottom two panels (IP:V5) show blots on material precipitated by anti-V5 beads. (C-J) Images of S2 cells transfected with pAw-Gal4 plasmid and *UAS-3xFLAG:Atro* (C), *UAS-vg:V5* (D), *UAS-sd:myc* (E), *UAS-3xFLAG:Atro* and *UAS-vg:V5* (F), *UAS-3xFLAG:Atro* and *UAS-sd:myc* (G), *UAS-vg:V5* and *UAS-sd:myc* (H), *UAS-3XFLAG:Atro*, *UAS-vg:V5*, and 300 ng of *UAS-sd:myc* (I), *UAS-3XFLAG:Atro*, *UAS-vg:V5*, and 37.5 ng of *UAS-sd:myc* (J), stained with antibodies to detect transfected proteins and Hoechst to label DNA.

To complement these co-immunoprecipitation experiments, we also examined potential co-localization amongst these proteins when expressed in S2 cells. On its own, Atro localized to the nucleus, where it concentrated into puncta, whereas Vg localized to both cytoplasm and nucleus, concentrating into puncta that generally appear smaller than Atro puncta (Fig. 7C,D). Sd staining exhibited a diffuse nuclear localization (Fig. 7E). When co-expressed together in S2 cells, Atro and Vg largely colocalized in puncta, mostly inside the nucleus, but with some protein also co-localized in the cytoplasm (Fig. 7F). This co-localization is consistent with the co-immunoprecipitation that we detected, further supporting the conclusion that they can associate together in the same complex.

Conversely, Atro and Sd did not co-localize, but instead largely appeared in mutually exclusive concentrations within the nucleus (Fig. 7G). Large Atro concentrations also visibly excluded DNA, whereas the Sd distribution was similar to the DNA distribution (Fig. 7C,G). Sd and Vg colocalized in large puncta within the nucleus, which overlap with DNA (Fig. 7H). When Sd, Atro, and Vg were co-expressed, all three proteins exhibited a dispersed distribution within the nucleus (Fig. 7I). As the Sd expression level was lowered, some colocalization of Atro and Vg in puncta could again be observed (Fig. 7J). These observations are consistent with the conclusion from co-immunoprecipitation experiments that Sd disrupts association between Vg and Atro.

### Atro associates with Vg through different regions than Sd does

To further investigate the association between Vg and Atro, we identified regions of Vg that are required for association with Atro in co-immunoprecipitation experiments. As it has previously been reported that Vg binds to Sd through a small central sequence motif (Sd-binding, amino acids 279-335) (Simmonds et al., 1998; Vaudin et al., 1999), we examined Atro association with this Vg-Sd-binding region, as well as Vg sequences N-terminal or C-terminal to this (Fig. S4B). We found that Atro co-precipitates the C-terminal fragment of Vg (Vg-C, amino acids 336-453) and more weakly associates with an N-terminal fragment (Vg-N, amino acids 1-278) but does not detectably co-precipitate the Vg-Sd-binding region (Fig. S4A). Within S2 cells, Atro accumulations were adjacent or encircled by Vg-N inside nuclei (Fig. S4C), Vg-N also made aggregates in the cytoplasm without Atro. Vg-Sd-binding region did not co-localize with Atro and accumulated as clumps in the cytoplasm, while Atro localized exclusively in the nuclei (Fig. S4D). Vg-C and Atro puncta overlap inside nuclei confirming their physical association (Fig. S4E). Thus, Atro and Sd do not directly compete for binding to the same region of Vg.

### Atrophin cooperates with Scribbler to regulate Hippo signaling in wing discs

Atrophin has previously been reported to act together with the corepressor Scribbler (Sbb), also known as Brakeless or Master of thickveins (Wehn and Campbell, 2006). To gain further insight into regulation of Hippo signaling by *Atro*, we investigated whether *sbb* has similar effects. Indeed, knockdown of *sbb* in posterior wing disc cells by RNAi led to upregulation of *ex-lacZ* near the DV boundary, and downregulation in proximal regions, similar to the effects of *Atro RNAi* (Fig. S5A). We also observed similar effects on expression of Diap1 (Fig. S5B). *sbb RNAi* was also associated with a reduction in nuclear Yki throughout the wing pouch, similar to *Atro RNAi* (Fig. S5C). As for Atro, this reduction in nuclear Yki could be explained by increased levels of Ds and a reduction in junctional levels of Dachs (Fig. S5D-E).

## Discussion

Our studies have identified multiple roles for Atro in modulating transcriptional regulatory networks within the developing *Drosophila* wing to direct Hippo signaling outputs (Fig. 8). Our observations reveal new layers of complexity in Hippo pathway regulation and emphasize how cell signaling outputs depend on interconnections with diverse transcriptional programs. In the case of Atro, these connections result in distinct effects on the expression of Hippo pathway target genes in different regions of the wing disc, such that when Atro is downregulated, expression of Yki target genes is elevated near the DV boundary but reduced in peripheral regions.

**Figure 8.**
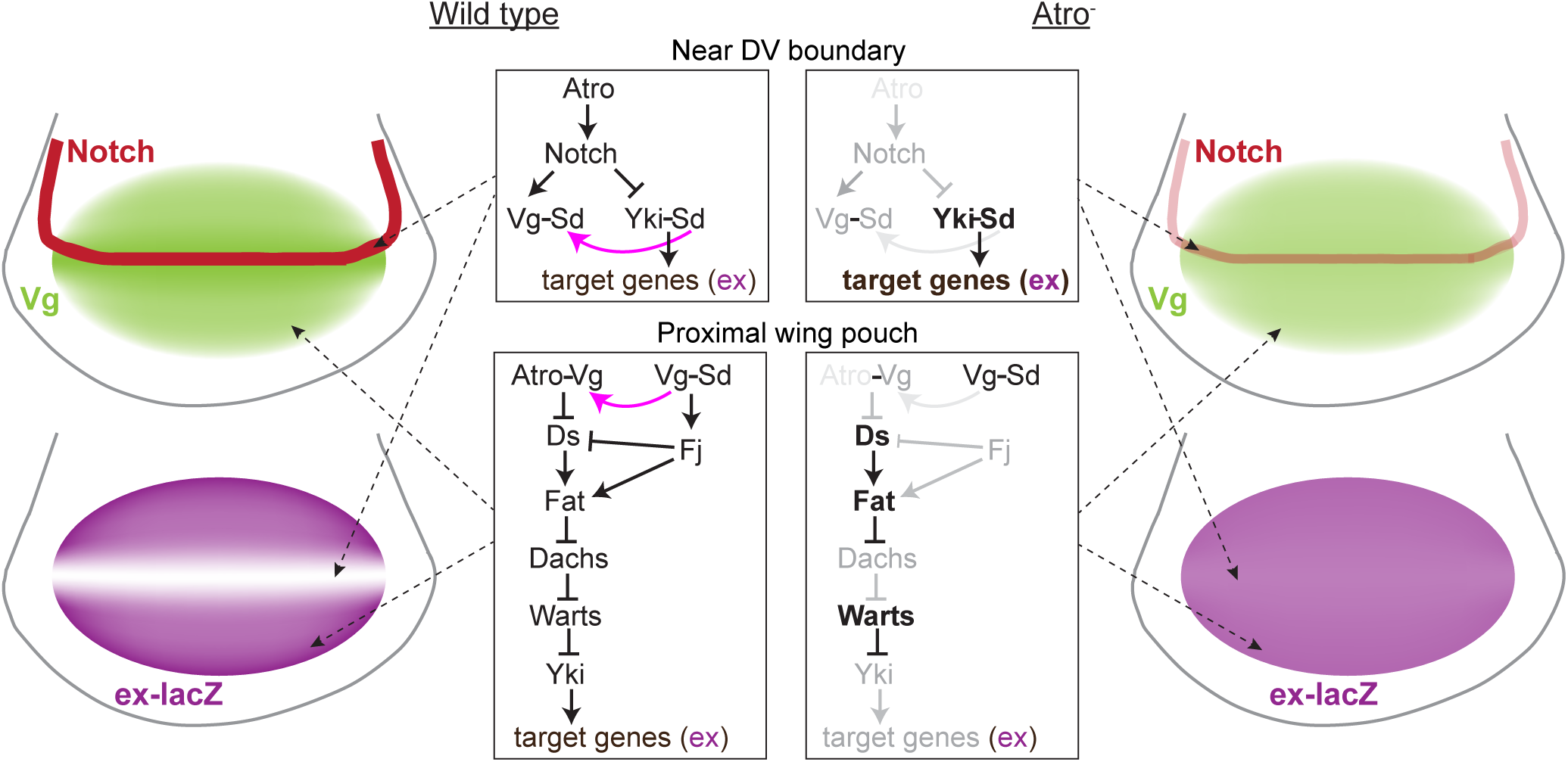
Summary Model of Atro regulation of Yorkie target genes. Schematic summary of Atro regulatory interactions modulating expression of Hippo pathway target genes in the developing wing. At right, in wildtype, Atro promotes Notch activation at DV boundary, which promotes expression of Vg. This suppresses Yki-Sd activity by pulling Sd away from Yki-Sd complexes into Vg-Sd complexes (purple arrow). Atro also enhances Yki activity by cooperating with Vg to repress Ds expression, which leads to lower levels of Fat activity, higher levels of Dachs at apical junctions, lower levels of Wts, and ultimately higher Yki activity. When Atro is knocked down, Notch activity is reduced, leading to lower Vg expression and consequently higher Yki activity near the DV boundary. Elsewhere, reduced Atro leads to elevated Ds, which increases Fat activity leading to lower Dachs, less inhibition of Wts, and consequently reduced Yki activity.

As Atro is a transcriptional co-repressor, the increased expression of Yki target genes near the D-V boundary upon loss of Atro could theoretically be consistent with Atro directly repressing expression of these genes. However, we favor instead the hypothesis that the Atro’s effects on Yki target genes are mediated through regulation of Notch and Vg. First, consistent with prior studies (Wang et al., 2018; Yeung et al., 2017), we observed that downregulation of Atro led to reduced expression of Notch target genes along the DV boundary of the wing disc, including Vg. The mechanism for this is not fully understood, although mis-regulation of fringe, a glycosyltransferase that modulates Notch signaling (Haines and Irvine, 2003), has been identified as a one possible factor (Yeung et al., 2017). An alternative explanation is suggested by studies in vertebrates, where the Atro homologue RERE was found to increase expression of Notch targets by stabilizing the cleaved Notch intracellular domain that mediates transcriptional activation (Wang et al., 2017). Regardless of the mechanism by which Notch signaling is impaired, prior studies have reported that Notch inhibits Yki activity along the DV boundary of wing discs (Djiane et al., 2014; Wang and Baker, 2018), at least in part through regulation of Vg, which competes with Yki for binding to Sd. This mechanism can account for our observations of increased Yki target gene expression when Atro is knocked down, and moreover explains why this effect is only observed near the DV boundary, where Notch is normally active, and why it is not associated with increased nuclear Yki and appears genetically to act in parallel with Yki regulation of target genes.

In contrast to the upregulation of Yki target genes near the D-V boundary when Atro is knocked down, we found that in more proximal regions of the wing Yki target genes were downregulated, and this was matched with a decrease in nuclear Yki. The decrease in Yki levels implies that Hippo signaling is reduced, and consistent with this we found lower levels of Dachs, which mediates one of the key upstream regulatory inputs into Hippo signaling in the wing: Dachs levels on membranes are downregulated by Ds-Fat signaling (Brittle et al., 2012; Mao et al., 2006), and Dachs on membranes, together with Dlish, inhibits Wts and Ex (Cho et al., 2006; Feng and Irvine, 2007; Vrabioiu and Struhl, 2015; Wang et al., 2019). The reduction in Dachs that we observe is likely due to the elevated Ds, and decreased Fj, that occur when Atro is knocked down. Less clear at present is how over-expression of Atro increases nuclear Yki and expression of Yki target genes in proximal regions, as we did not observe statistically significant increases in Dachs levels.

The upregulation of Ds and downregulation of Fj when Atro is knocked down are reminiscent of the effects of Vg on Ds and Fj (Cho and Irvine, 2004; Zecca and Struhl, 2010), which led us to further investigate functional and physical connections between them. We found that knockdown of Atro was epistatic to Vg over-expression for regulation of Ds expression. This indicates that the effect of Atro cannot be explained by downregulation of Vg and is instead consistent with the possibility that Vg requires Atro to repress Ds. We further found that Atro can physically associate with Vg, which supports a model in which Atro and Vg act together to repress Ds. Since we did not detect a trimeric Atro-Vg-Sd complex, and instead found that Atro and Sd compete for association with Vg, we infer that Atro pulls Vg away from a transcriptional activation complex with Sd to instead participate in a distinct transcriptional repression complex with Atro, which downregulates Ds. The mechanism of Atro-Sd competition appears to be indirect, as they each associate with distinct regions of Vg. As neither Atro nor Vg are DNA binding proteins, we infer that an Atro-Vg complex would need to associate with other factors to effect repression of Ds. One such factor identified by our studies, though also not a DNA binding protein, is the Atro co-factor Sbb. In addition to upregulation of Ds, knock down of Atro also decreased expression of *fj*, and multiple factors that could contribute to this, including decreased Vg expression, removal of Vg from Vg-Sd complexes, and decreased Yki activity, as Fj is a downstream target of both Sd-Vg and Sd-Yki. Altogether, these observations enhance our understanding of Hippo pathway regulation and identify new roles for Atro during wing development.

## Materials and Methods

### Plasmids and cloning

An expression plasmid for *Atro* was created by amplifying a full length *Atro* cDNA (Atro-RA variant) from plasmid pPFW (Yeung et al., 2017). As DNA sequencing identified three differences (T93P, D103G, Y902C) compared to the sequence in Flybase they were corrected using a Q5 Site-Directed Mutagenesis Kit (NEB #E0552S) according to the manufacturers protocol. DNA fragments were then PCR amplified and inserted into pUAST-3XFLAG vector using NEBuilder HiFi DNA Assembly Master Mix (NEBuilder E2621L) to generate pUAST-3xFLAG:Atro. A V5-tagged *vg* expression plasmid was constructed by PCR amplifying the *vg* coding region from *UAS-vg [49]* (Kim et al., 1996) and inserting it first into pAc5.1-V5:6XHis to generate pAc5.1-vg:V5:6xHis and then PCR amplifying vg:V5:6XHis and inserting into pUAST to generate pUAST-vg:V5:6XHis. Myc tagged *vg* and *sd* expression plasmids pUAST-vg:Myc and pUAST-sd:Myc were constructed through PCR amplification of *vg* and *sd* with a Myc tag sequence added to the PCR primer, and then inserting the amplified DNA into pUAST. The *vg* fragments *vg-N* (1-278), vg-Sd-binding (279-335), and vg-C (336-453) were fused with a C terminal GFP:V5:His tag by PCR amplifying vg fragments and GFP:V5:His and inserting them into pUAST to generate pUAST-vg-N:GFP:V5:His, pUAST-vg-Sd-binding:GFP:V5:His, and pUAST-vg-C:GFP:V5:His.

### Drosophila genetics

Transgenic flies expressing UAS-3xFLAG:Atro inserted at 28E7 were created by Bestgene after injection of pUAST-3XFLAG:Atro plasmid. A UAS-Atro transgene inserted on the third chromosome was a gift from Antonio Baonza (Charroux et al., 2006).

For temperature shift experiments, flies were cultured at 18 °C for 5-6 days and then transferred to 29° C to induce transgene expression for 48 h in RNAi knock down experiments and for 24 h in over-expression experiments. RNAi experiments included co-expression of Dcr2 under UAS-Dcr2 control to enhance the efficiency of RNAi (Dietzl et al., 2007), and which with anti-Dcr2 antibody staining can also serve as a marker for Gal4 expressing cells.

Gal4-expressing clones were generated in flies expressing the FLP-out cassette *act>CD2>Gal4* by heat shocking larvae at 60-72h AEL for 10 min at 38 °C and marked by expression of UAS-RFP. After heat shock larvae were maintained at 25 °C for 48 h before dissection.

RNAi lines used were:

*UAS-RNAi-Atro[HMS00756]* (FBsf0000169110)

*UAS-RNAi-Atro[KK100675]* (FBsf0000086814)

*UAS-RNAi-yki[KK109756]* (FBsf0000092436)

*UAS-RNAi-yki[HMS00041]* (FBsf0000382991)

*UAS-RNAi-sbb [JF02375]* (FBsf0000086425) (Perkins et al., 2015)

Reporter lines used were:

*ex-lacZ* (Boedigheimer et al., 1997)

*vg[BE]-lacZ* (Williams et al., 1994)

*PBac[gug-GFP.FPTB}* (FBal0356988)

*fj-lacZ[P1]* (FBal0049503) (Brodsky and Steller, 1996)

*wts-lacZ^P2^* (FBal0044496)

*kibra-lacZ* (FBst0011567)

*jub:GFP* (Sabino et al., 2011)

*dachsous:GFP* (Brittle et al., 2012)

PBac{dachs.GFP} (FBal0269880) (Bosveld et al., 2012)

Over-expression lines used were:

UAS-vg [49] (Kim et al., 1996)

*UAS-ds(III)* (Matakatsu and Blair, 2004)

*UAS-yki:V5^S168A^* (Oh and Irvine, 2008)

*UAS-3xFLAG:Atro* (this work)

Gal4 lines used were:

*en-Gal4* (FBti0003572) (Perrimon Lab)

*tub-Gal80^ts^* (McGuire et al., 2004)

act>CD2>Gal4 (Pignoni and Zipursky, 1997)

hh-Gal4 (FBti0017278)

### Immunostaining wing discs

As described previously (Rauskolb and Irvine, 2019), mid to late third instar larvae were dissected in Ringer’s solution (111 mM NaCl, 1.88 mM KCl, 2.38 mM NaHCO_3_, 0.0641 mM NaH_2_PO_4_-2H_2_O, 1.081 mM CaCl_2_) and fixed in 4% paraformaldehyde in PBS (155mM NaCl, 1mM KH2PO4, 3mM Na2HPO4, pH 7.4) for 20 min (or 12 min for Dachs:GFP and Dachsous:GFP discs), rinsed twice in Ringer’s solution and PBT (PBS, 1% BSA, and 0.1% Triton X-100), and washed twice (20 min each) in PBT. Larvae were then blocked for 30 min in PBT with 5% donkey serum and incubated in primary antibodies overnight (in PBT+5% donkey serum) at 4 °C with gentle rocking. Larvae were then washed 4 times 10 min each in PBT, blocked in PBT with 5% donkey serum, and incubated with secondary antibodies. After incubation, larvae were washed 2x in PBT, stained with Hoechst 33342 (Invitrogen, H3570) for 20 min and then washed again 2x in PBT. Wing discs were dissected and mounted on the slides using Vectashield Antifade Mounting Medium (Vector Laboratories, H-1000-10).

All antibodies used in this study are listed in Supplementary Table 1.

**Table 1.** Antibodies used.

| Reagent | Source | Identifier |
| --- | --- | --- |
| Anti-Yki (Rabbit) (1:80) | Irvine Lab (Oh and Irvine, 2008) | N/A |
| Anti-βGal (Mouse) (1:200) | Developmental Studies Hybridoma Bank | JIE7 |
| Anti-βGal (Chicken) (1:200) | Abcam | ab9361 |
| Anti-Dcr2 (Rabbit) (1:800) | Abcam | ab4732 |
| Anti-Diap1 (Rabbit) (1:200) | Bruce Hay | N/A |
| Anti-Notch, intracellular domain (Mouse) | DSHB | C17.9c6 |
| Anti-Vg (Rabbit) (1:100) | K. Guss (Guss Lab) | N/A |
| Anti-Atro (Rabbit) (1:500) | H. McNeill (McNeill Lab)(Yeung et al., 2017) | N/A |
| Anti-Atro (Mouse) (1:20) | Mattias Mannervik (Yeung et al., 2017) | N/A |
| Anti-E-cadherin (Rat) (1:100) | Developmental Studies Hybridoma Bank | DCAD2-c |
| Anti-Engrailed (Mouse) (1:100) | DSHB | 4D9 |
| Anti-FLAG (Mouse) (1:500) (IF S2 cells) | Sigma | Cat#F3165; RRID:AB_259529 |
| Anti-FLAG (Mouse) (1:200) (IF tissue) | Sigma | Cat#F3165; RRID:AB_259529 |
| Anti-FLAG (Mouse) (1:2000) (Western) | Sigma | Cat#F3165; RRID:AB_259529 |
| Anti-FLAG (Rabbit) (1:500) (IF S2 cells) | Sigma | Cat#F7425; RRID:AB_439687 |
| Anti-FLAG (Rabbit) (1:200) (IF tissue) | Sigma | Cat#F7425; RRID:AB_439687 |
| Anti-FLAG (Rabbit) (1:2000) (Western) | Sigma | Cat#F7425; RRID:AB_439687 |
| Anti-V5 (Mouse) (1:500) (IF S2 cells) | Invitrogen | Cat#R960-25; RRID:AB_2556564 |
| Anti-V5 (Mouse) (1:200) (IF tissue) | Invitrogen | Cat#R960-25; RRID:AB_2556564 |
| Anti-V5 (Mouse) (1:5000) (Western) | Invitrogen | Cat#R960-25; RRID:AB_2556564 |
| Anti-V5 (Rabbit) (1:500) (IF S2 cells) | Bethyl Laboratories | Cat#A190-120A; RRID:AB_67586 |
| Anti-V5 (Rabbit) (1:200) (IF tissue) | Bethyl Laboratories | Cat#A190-120A; RRID:AB_67586 |
| Anti-V5 (Rabbit) (1:5000) (Western) | Bethyl Laboratories | Cat#A190-120A; RRID:AB_67586 |
| Anti-c-myc (Rat) (1:200) (IF S2 cells) | BIO-RAD | Cat#MCA1929; RRID:AB_322203 |
| Anti-c-myc (Rat) (1:1000) (Western) | BIO-RAD | Cat#MCA1929; RRID:AB_322203 |
| Anti-myc (Mouse) (1:1000) (Western) | Invitrogen | Cat#MA1-980<br>RRID:AB_558470 |
| Anti-GAPDH (Mouse) (1:20,000) (Western) | Proteintech | Cat#60004-1-IG<br>RRID:AB_2107436 |
| Donkey anti-Mouse IgG (H+L) Alexa Fluor™ 488 (1:100 IF tissue, 1:500 IF S2 cells) | Invitrogen | Cat#A-21202;<br>RRID:AB_141607 |
| Donkey anti-Rabbit IgG (H+L) Alexa Fluor™ 488 (1:100 IF tissue, 1:500 IF S2 cells) | Invitrogen | Cat#A-21206;<br>RRID:AB_2535792 |
| Donkey anti-Chicken IgY (H+L) Alexa Fluor™ 488 (1:100 IF tissue) | Invitrogen | Cat#A78948;<br>RRID:AB_2921070 |
| Cy-3AffiniPure Donkey anti-Rabbit IgG (H+L) | Jackson Immuno Research Labs | Cat#711-165-152;<br>RRID:AB_2307443 |
| Cy3-AffiniPure Donkey Anti-Mouse IgG (H+L) | Jackson Immuno Research Labs | Cat#715-165-151;<br>RRID:AB_2315777 |
| Cy3-AffiniPure Donkey Anti-Rat | VWR | Cat#102649-822 |
| Donkey anti-Rabbit IgG (H+L) Alexa Fluor 647-AffiniPure (1:100 IF tissue, 1:500 IF S2 cells) | Jackson Immuno Research Labs | Cat#711-605-152;<br>RRID:AB_2492288 |
| Donkey anti-Mouse IgG (H+L) Alexa Fluor 647-AffiniPure | Jackson Immuno Research Labs | Cat#715-605-151;<br>RRID:AB_2340863 |
| Donkey-anti-Rat 647 (1:100 IF tissue, 1:500 IF S2 cells) | Invitrogen | Cat#715-165-153<br>RRID: AB_2340694 |
| IRDye 800CW Donkey anti-Rabbit IgG (1:20,000) (Western) | LICORbio | Cat#926-32213<br>RRID:AB_621848 |
| IRDye 680RD Donkey anti-Mouse IgG (1:20,000) (Western) | LICORbio | Cat#926-68072<br>RRID:AB_10953628 |

### S2 cells transfection, co-immunoprecipitation, western blotting, and immunostaining

*D. melanogaster* S2 cells were cultured at 28 °C in Schneider’s *Drosophila* Medium (Gibco, 21720001), supplemented with 10% fetal bovine serum (ATCC, 30-2020) and antibiotics (Gibco, 15240062). Cells were transfected in six-well plates with plasmids containing the UAS promoter induced by added pAw-Gal4 vector (Seth Blair) to express Gal4 under *act5c* promoter. The following vectors were used: pUAST-attB, pUAST-3XFLAG:Atro, pUAST-vg:V5:His, pUAST-sd:Myc, pAw-Gal4, pUAST-GFP:3XFLAG, pUAST-vg:Myc, pUAST-vg-N:GFP:V5:6xHis, pUAST-vg-Sd-binding:GFP:V5:6xHis, pUAST-vg-C:GFP:V5:6xHis. Transfections were carried out using the Effectene Transfection Reagent following the manufacturer’s recommendations (QIAGEN, 301427). Briefly, S2 cells were transfected with a combination of the above-mentioned plasmids for 48 h at 28 °C.

For immunoprecipitation and western blotting, cells were then collected and lysed in 400 µL of RIPA buffer (140 mM NaCl, 10 mM Tris-HCl pH 8.0, 1 mM EDTA pH 8.0, 1.0% Triton X-100, 0.1% SDS, and 0.1% sodium deoxycholate) supplemented with protease (Roche, 11873580001) and phosphatase (Millipore, 524625) inhibitors for 30 min. The lysate’s supernatant was divided in two: 20 µL served as input while the rest was diluted in 400 µL of dilution buffer (10 mM Tris-HCl pH 7.4, 0.5 mM EDTA, 150 mM NaCl) and pre-cleared with 25 μl of Pierce Protein A agarose beads for an hour (Thermo Fisher Scientific, 20333). Co-immunoprecipitation was then performed using 25 μl of either ChromoTek DYKDDDDK Fab-Trap™ Agarose (Proteintech, Va) or ChromoTek V5-Trap Magnetic Agarose (Proteintech, v5tma) beads for 1 hr. Samples were then washed three times with 800 μl of dilution buffer supplemented with protease and phosphatase inhibitors. The bead slurry and the inputs were resuspended in Laemmli buffer, denatured and loaded on 4-15 % gradient SDS-PAGE Mini-Protean TGX gel (Bio-Rad, 4561086). Proteins were then transferred to a nitrocellulose membrane (Bio-Rad, 170-4270) using Trans-Blot Turbo (BioRad). Membranes were blocked with Intercept Blocking Buffer (LI-COR, 927-70001) for 1 h and primary antibodies added for incubation overnight at 4 °C with gentle shaking. After being washed, membranes were probed with secondary antibodies for 2 h at room temperature, washed and imaged on Odyssey CLX Imaging System (LI-CORbio).

For immunostaining, coverslips were coated with ConcanavalinA (Millipore Sigma, L7647) for 1 h and dried for 2 h. 50 μl of transfected S2 cells were added to 450 μl of S2 cell media and mixture was transferred to the coated cover slips to attach for 2 h. The cells were fixed with 4% paraformaldehyde in PBS for 10 min and washed for 5 min in PBS. After washing, the cells were permeabilized for 10 min and blocked for 1 h in PBT (PBS, 1% BSA, and 0.1% Triton X-100) on a rocking platform. The cells were then incubated with primary antibodies in PBT supplemented with 5% donkey serum overnight at 4 °C. Cells were then washed three times with PBS and stained with secondary antibodies followed by staining with Hoechst 33342 (Invitrogen, H3570) for 10 min and then mounted on the slides using Dako Fluorescence Mounting Media (Agilent Technologies, S3023). Samples were imaged using a Leica SP8 confocal microscope.

### Image quantitation

Fluorescence signal intensities in wing discs were quantified using Fiji. For Yki and ex-lacZ analysis, nuclear signals in a single Z plane were determined by segmenting nuclei based on DNA stains, applying a median filter, and thresholding using Phansalkar or Default methods. Anterior and posterior ROIs of the same size were defined manually and ratios between ROIs at the same proximal-distal location compared. Dachs:GFP and Jub:GFP signal intensities were quantified using a previously described MATLAB script (Ibar et al., 2023).

### Statistical analysis

Statistical analysis was performed using GraphPad Prism, and Student’s *t-test* was used for pairwise comparisons. All quantifications are presented as means ± s.e.m. For all statistical tests *p* values are: *\*p < 0.05, **p < 0.001, ***p < 0.0001, ****p < 0.00001*.

## Acknowledgments

This research used antibodies obtained from the Developmental Studies Hybridoma Bank, fly stocks from the Bloomington *Drosophila* Stock Center, plasmids from *Drosophila* Genomics Resource Center, information from Flybase, and confocal microscopes at the Waksman Institute Shared Imaging Facility. We also thank Helen McNeill, Nick Baker, Antonio Baonza, Kirsten Guss, Seth Blair, Bruce Hay, Mattias Mannervik, for *Drosophila* stocks, plasmids and antibodies, and Derrick Michell and Mayank Chauhan for experimental assistance. This research was supported by National Institutes of Health grant GM131748 (KDI).

## Author Contributions

Conceptualization, Writing: DM, KDI; Methodology, Investigation: DM, CR, TL, SV, LC; Supervision, Funding Acquisition: KI.

## Supplemental Figure Legends

**Figure S1.**
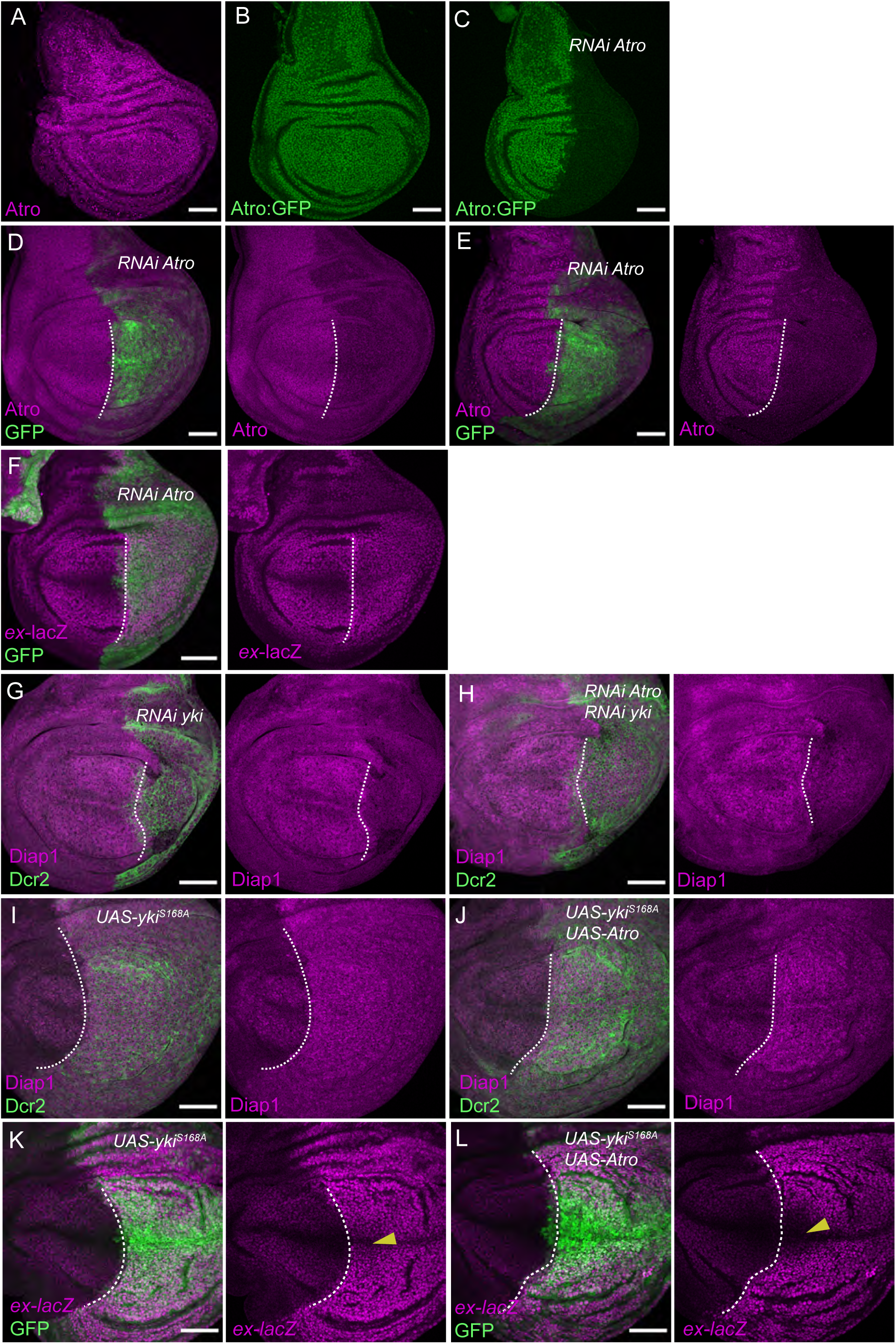
Atro knock down and effect on Yki target genes. (A) Expression of Atro protein in OR wing disc. (B) Expression of Atro:GFP from a genomic construct (PBac-Gug-GFP.FPTB). (C) Expression of Atro:GFP in wing discs expressing *en-Gal4 tub-Gal80^ts^ UAS-Dcr2* crossed to Atro:GFP *UAS-RNAi-Atro[HMS00756].* (D,E) Expression of Atro protein in wing discs expressing *en-Gal4 UAS-GFP ex-lacZ tub-Gal80^ts^ UAS-Dcr2* crossed to *UAS-RNAi-Atro[HMS00756]* (D) and *UAS-RNAi-Atro[KK100675]* (E), stained with mouse (D) or rabbit (E) anti-Atro. (F) Expression of *ex-lacZ* (magenta) in third instar wing discs from *en-Gal4 UAS-GFP ex-lacZ; tub-Gal80^ts^ UAS-Dcr2* crossed to *UAS-RNAi-Atro[KK100675]*. Posterior compartment is labeled by GFP expression (green). (G-J) Expression of Diap1 (magenta) in wing discs from *en-Gal4; tub-Gal80^ts^ UAS-Dcr2* crossed to *UAS-RNAi-yki [KK109756]* (G), *UAS-RNAi-yki [KK109756]* and *UAS-RNAi-Atro[HMS00756]* (H*), UAS-yki^S168A^* (I), *UAS-yki^S168A^* and *UAS-Atro* (J). Posterior compartment is labeled by Dcr2 (green). (K,L) Expression of *ex-lacZ* (magenta) in third instar wing discs from *en-Gal4 UAS-GFP ex-lacZ; tub-Gal80^ts^ UAS-Dcr2* crossed to *UAS-yki^S168A^* (K) or *UAS-yki^S168A^* and *UAS-3xFLAG:Atro* (L). Posterior compartment is labeled by GFP expression (green). Dashed lines mark anterior-posterior compartment boundary, yellow arrowheads point to the DV boundary region, Scale bars=50 μm.

**Figure S2.**
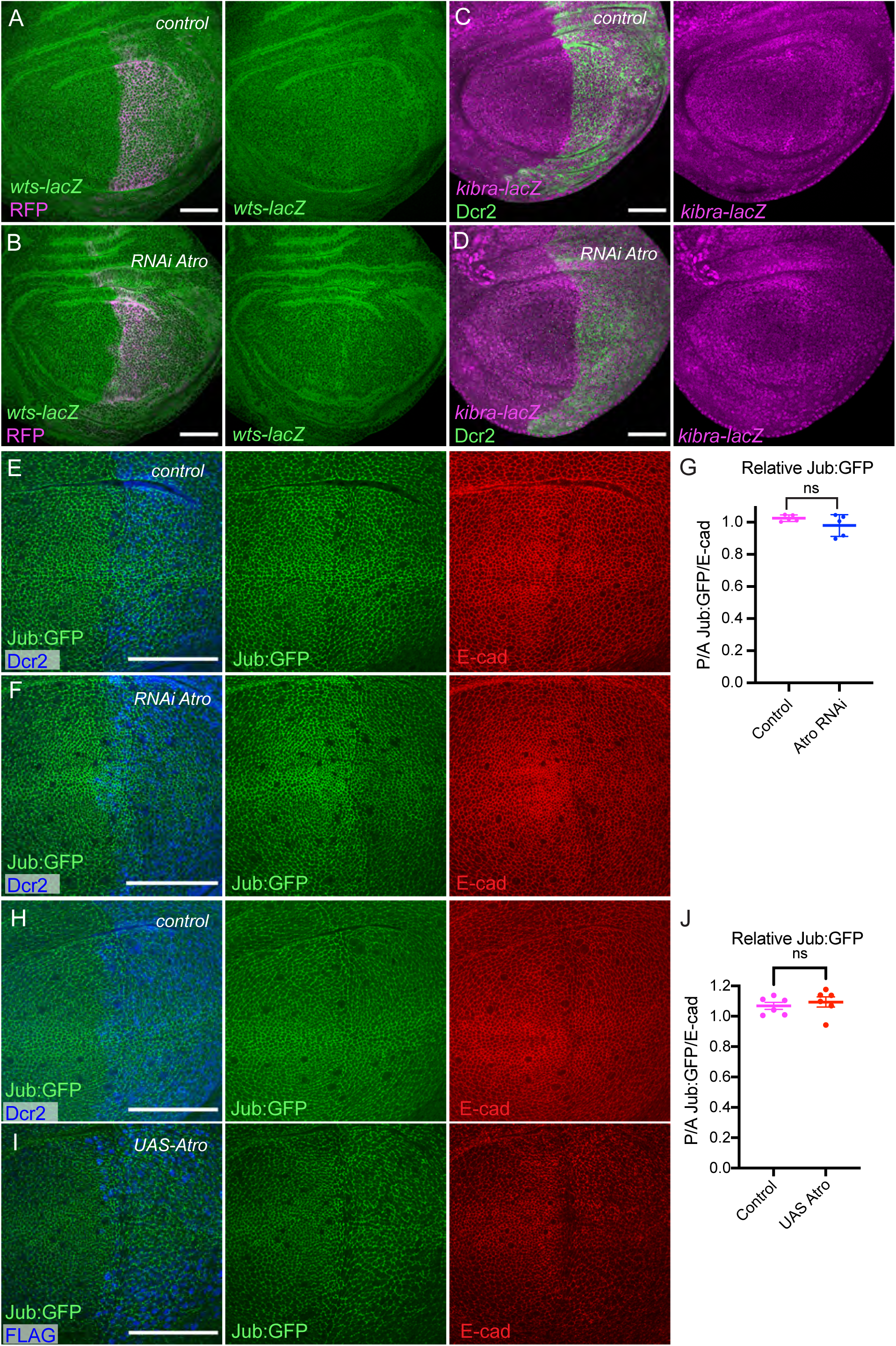
Atro regulation of Yki is not explained by regulation of Wts, Kibra, or Jub. (A-B) *wts-lacZ* expression (green) in wing discs from *en-Gal4 UAS-RFP tub-Gal80^ts^ UAS-Dcr2* crossed to control (OR) (A) or *UAS-RNAi-Atro[KK100675]*. Posterior compartment is labeled by RFP (magenta). (C-D) *kibra-lacZ* expression (magenta) in wing discs from *hh-Gal4 tub-Gal80^ts^ UAS-Dcr2* crossed to control (OR) (C) or *UAS-RNAi-Atro[KK100675].* Posterior compartment is labeled by Dcr2 (green). (E,F,H,I) Jub:GFP (green) in wing discs expressing *en-Gal4 tub-Gal80^ts^ jub:GFP UAS-Dcr2* crossed to control (OR) (E), *UAS-RNAi-Atro[HMS00756]* (F), control (OR) (H), or UAS-3xFLAG:Atro (I). Posterior compartment is labeled by Dcr2 (blue) or FLAG:Atro (blue). Scale bars=50 μm. (G) Quantitation of relative Jub:GFP/E-cad in posterior versus anterior cells from wing discs of the genotypes in E,F. Significance of differences calculated by t test, N=5(control), N=8(Atro RNAi). (J) Quantitation of relative Jub:GFP/E-cad in posterior versus anterior cells from wing discs of the genotypes in H,I. Significance of differences calculated by t test, N=6(control), N=6(UAS-Atro).

**Figure S3.**
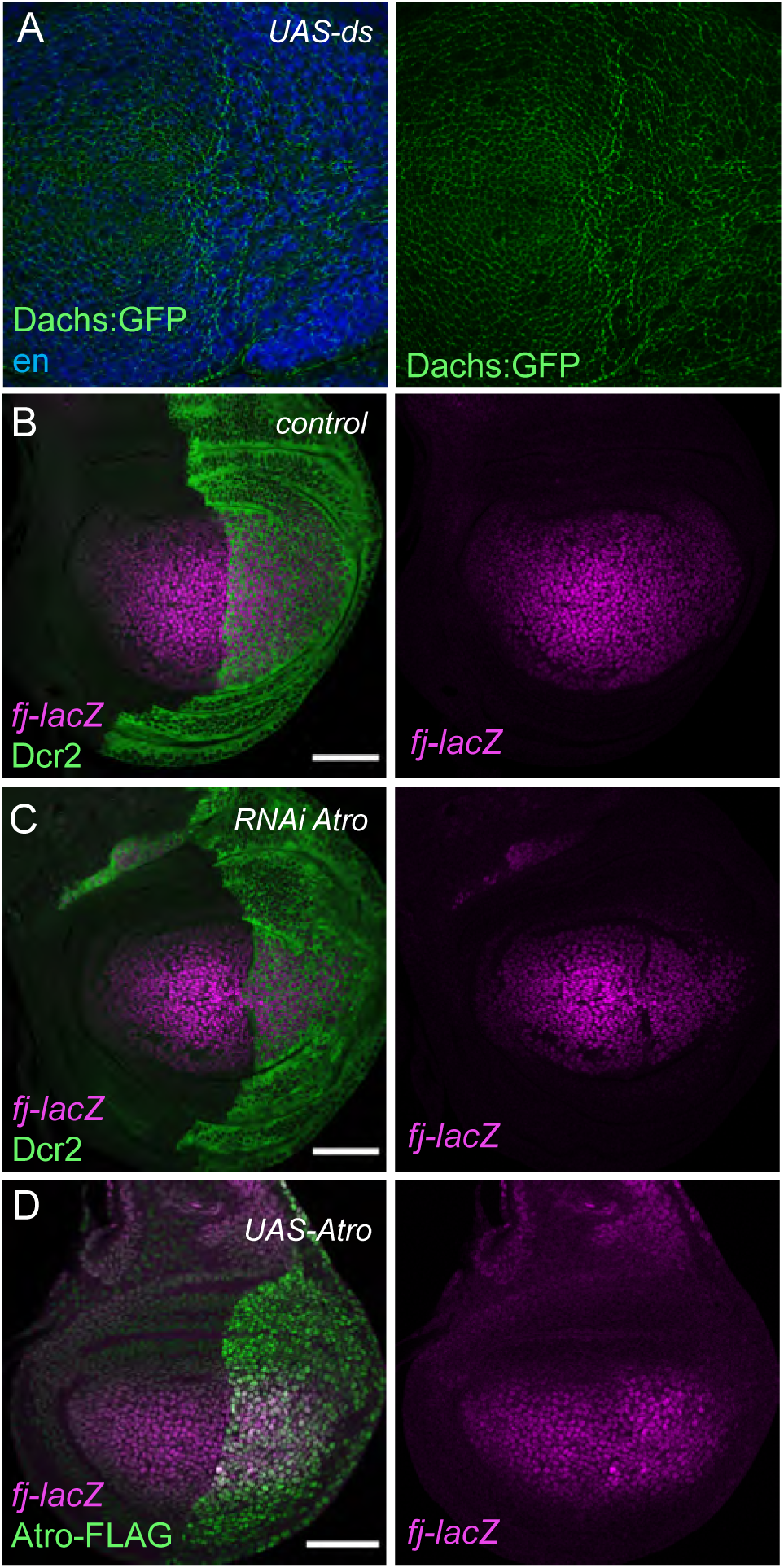
Additional analysis of Atro effects on Ds-Fat pathway components. (A) Dachs:GFP in wing discs expressing *en-Gal4 tub-Gal80^ts^ Dachs:GFP* crossed to *UAS-ds*. Posterior compartment is labeled by engrailed (en) staining (blue). (B-D) Expression of *fj-lacZ* (magenta) in wing discs expressing *fj-lacZ* hh*-Gal4 tub-Gal80^ts^ UAS-Dcr2* crossed to control (OR) (B)*, UAS-RNAi-Atro[HMS00756]* (C), or *UAS-3xFLAG:Atro* (D). Posterior compartment is labeled by Dcr2 or Atro:FLAG (green).

**Figure S4.**
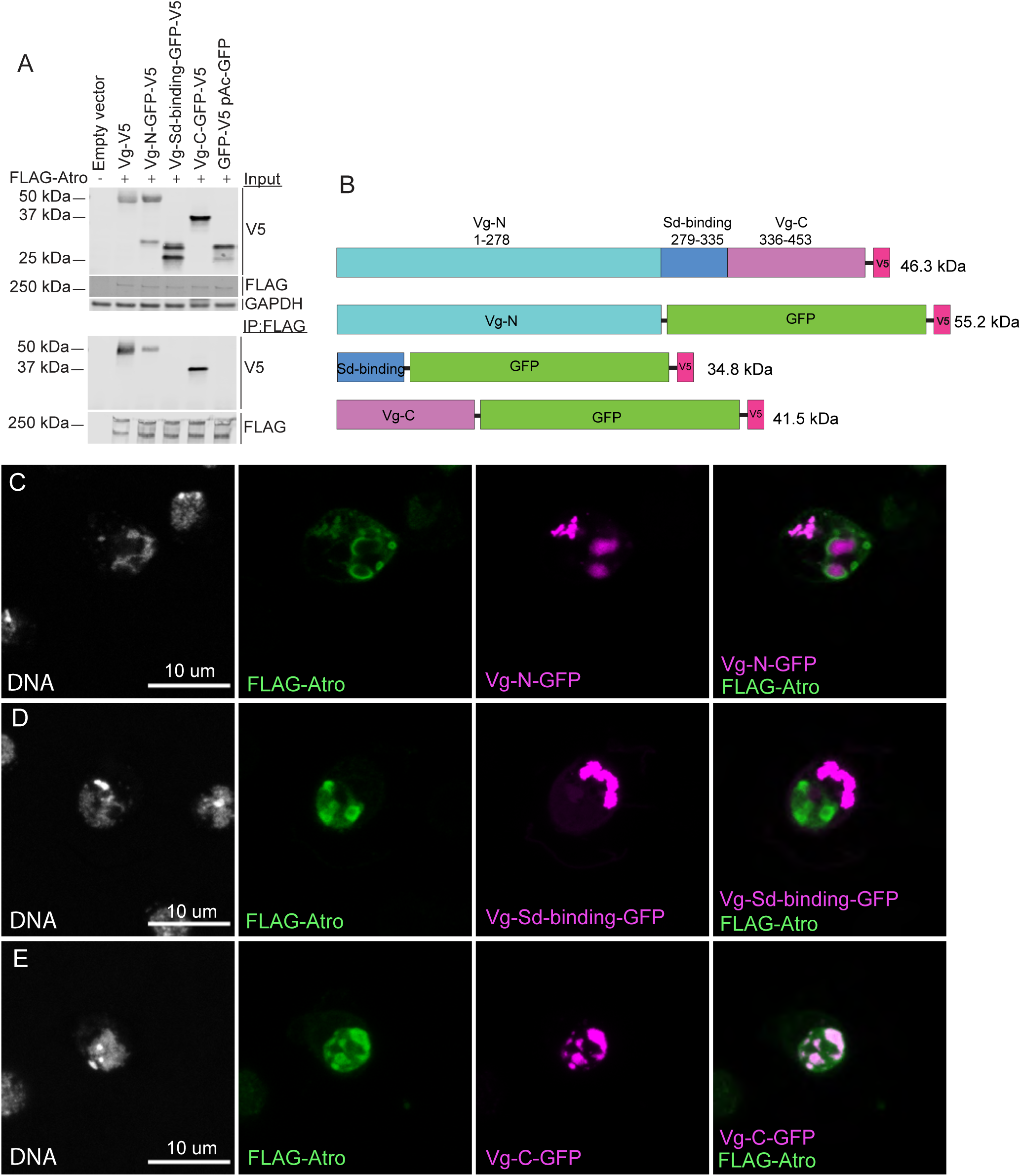
Atro-Vestigial interaction maps to N- and C-terminal regions. (A) Western blots showing results of co-immunoprecipitation experiments on lysates of S2 cells transfected with plasmids expressing the proteins indicated at top. Antibodies used for detection indicated at right of blots, location of protein size standards indicated at left. Top 3 panels (Input) show blots on cell lysates, bottom 2 panels (IP:FLAG) show blots on proteins precipitated by anti-FLAG beads. (B) Schematic representation of Vestigial protein domains and protein fragments used to map association with Atro. (C-E) Images of S2 cells transfected with act-Gal4 plasmid and *UAS-3xFLAG:Atro* and UAS-vg-N:V5:6xHis (C), *UAS-3xFLAG:Atro* and UAS-vg-Sd-binding:V5:6xHis (D), *UAS-3xFLAG:Atro* and UAS-vg-C:V5:6xHis (E), stained with antibodies to detect transfected proteins and Hoechst to label DNA.

**Figure S5.**
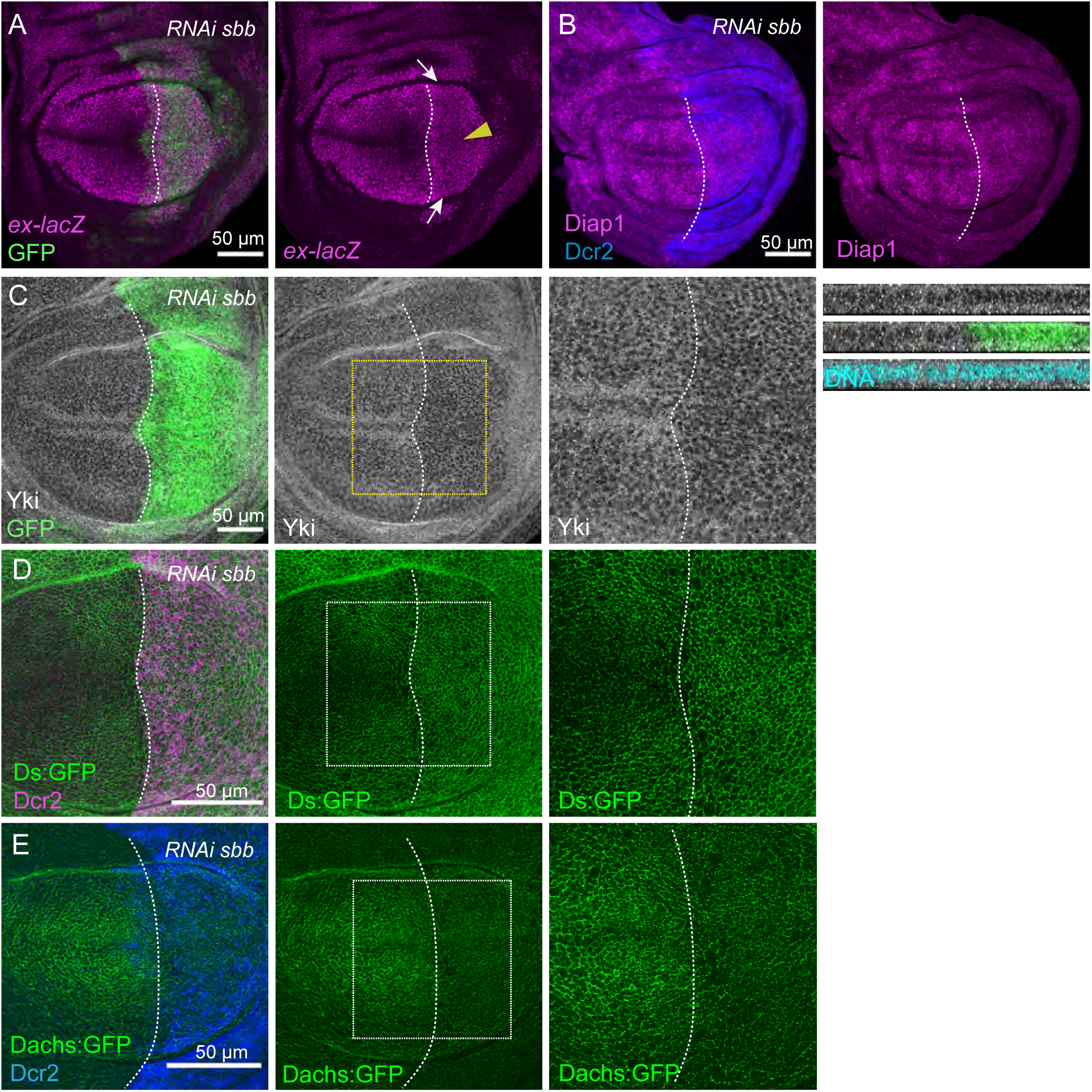
Scribbler influences Hippo signaling similarly to Atro. (A) Expression of *ex-lacZ* (magenta) in third instar wing discs from *en-Gal4 UAS-GFP ex-lacZ tub-Gal80^ts^ UAS-Dcr2* crossed to *UAS-RNAi-sbb*. Posterior compartment is labeled by GFP (green). (B) Expression of Diap1 (magenta) in third instar wing discs from *en-Gal4 tub-Gal80^ts^ UAS-Dcr2 Dachs:GFP* crossed to *UAS-RNAi-sbb*. Posterior compartment is labeled by Dcr2 (blue). (C) Yki protein staining (white) in wing discs expressing *en-Gal4 UAS-GFP ex-lacZ tub-Gal80^ts^ UAS-Dcr2* crossed to *UAS-RNAi-sbb*, with posterior compartment marked by expression of GFP (green). Yki stain alone in horizontal section is shown to the right, with dashed yellow box outlining region shown at higher magnification in the panels further to the right. Images right of this show vertical sections through the discs, including Yki, Yki and GFP, Yki and a DNA stain as a nuclear marker (cyan). (D) Ds:GFP in wing disc expressing *hh-Gal4 tub-Gal80^ts^ UAS-Dcr2 dachsous:GFP* crossed to *UAS-RNAi-sbb*, with posterior compartment labeled by Dcr2 (magenta). Panels to right show Ds:GFP alone, and a higher magnification image of Ds:GFP. (E) Dachs:GFP (green) in wing disc expressing *en-Gal4 tub-Gal80^ts^ UAS-Dcr2 Dachs:GFP* crossed to *UAS-RNAi-sbb*, with posterior compartment labeled by Dcr2 (blue). Panels to right show Dachs:GFP alone, and a higher magnification image of Dachs:GFP. Scale bar=50µm.

